# Structural Basis of GPCR-G Protein Pre-coupling and Activation: Insights from CCR1-Gi Complex

**DOI:** 10.1101/2024.11.01.621549

**Authors:** Zhehua Shao, Yingjun Dong, Ruixin Jia, Qingya Shen, Bingpeng Yao, Xinheng He, Qingning Yuan, Dandan Shen, Chunyou Mao, Chao Zhang, Zhihua Chen, H. Eric Xu, Songmin Ying, Yan Zhang, Wen Li

## Abstract

GPCR-mediated G protein activation cycle contains obligated step of the receptor-G protein assembly before G protein activation (pre-coupling state). However, transient nature of this pre-coupling state has prevented structural and mechanistic understanding of this essential step of G protein activation cycle. In this study, we discover that CCR1 has high level of pre-coupling state, which allows its rapid response to chemokines. Taking advantage of this observation, we uncover molecular mechanism of the pre-coupling state by solving the cryo-electron microscopy structure of the chemokine receptor CCR1 in complex with its cognate G protein (Gi) in a pre-coupled, inactivated state. This structure reveals that CCR1 adopts a conformation distinct from both its fully active and inactive states, with the G protein’s α5 helix partially inserted into the receptor’s intracellular cavity. Notably, the C-terminal four residues of the Gα subunit are disordered in this pre-coupled state, contrasting with their well-defined α-helical structure in the fully active complex. Functional assays demonstrate that while deletion of these four C-terminal Gα residues does not affect pre-coupling, it abolishes G protein activation upon agonist binding. This finding highlights the critical role of these residues in GPCR-mediated G protein activation, but not in initial recruitment. Furthermore, our studies indicate that the ability to form pre-coupled complexes is conserved among chemokine receptors, suggesting a common mechanism for rapid signal transduction in this GPCR subfamily. These results provide the first structural evidence for GPCR-G protein pre-coupling and offer molecular insights into the transition from inactive to active states. Our findings fill the long-missing gap in understanding GPCR-mediated G protein activated cycle.

## Introduction

G protein-coupled receptors (GPCRs) constitute the largest family of membrane proteins in eukaryotes, playing pivotal roles in diverse physiological processes and serving as targets for approximately 35% of all FDA-approved drugs^1,2^. These receptors transduce extracellular signals across the plasma membrane by coupling to heterotrimeric G proteins, initiating intracellular signaling cascades that regulate cellular responses^3^. While the canonical model of GPCR activation posits that agonist binding induces conformational changes in the receptor, promoting its interaction with the C-terminal α5 helix of the Gα subunit protein, which induces subsequent activation of G proteins. However, accumulating evidence suggests a more nuanced picture of the process of ligand-bound GPCR-mediated G protein activation^4,5^.

Numerous functional and computational studies have indicated that some GPCRs can form complexes with their cognate G proteins prior to agonist binding, a phenomenon termed “pre-coupling” (Figure 1a). In this pre-coupling state, neither receptor nor G protein is in the activated state, but their association could enable rapid signal transduction upon ligand binding, allowing for swift cellular responses to external stimuli^6–12^. However, despite the potential physiological importance of pre-coupling, structural evidence for these complexes has remained elusive. The lack of high-resolution structures has hindered our understanding of the molecular determinants governing GPCR-G protein pre-coupling and the conformational changes that occur during the transition from pre-coupled to fully active states.

**Figure 1|.**
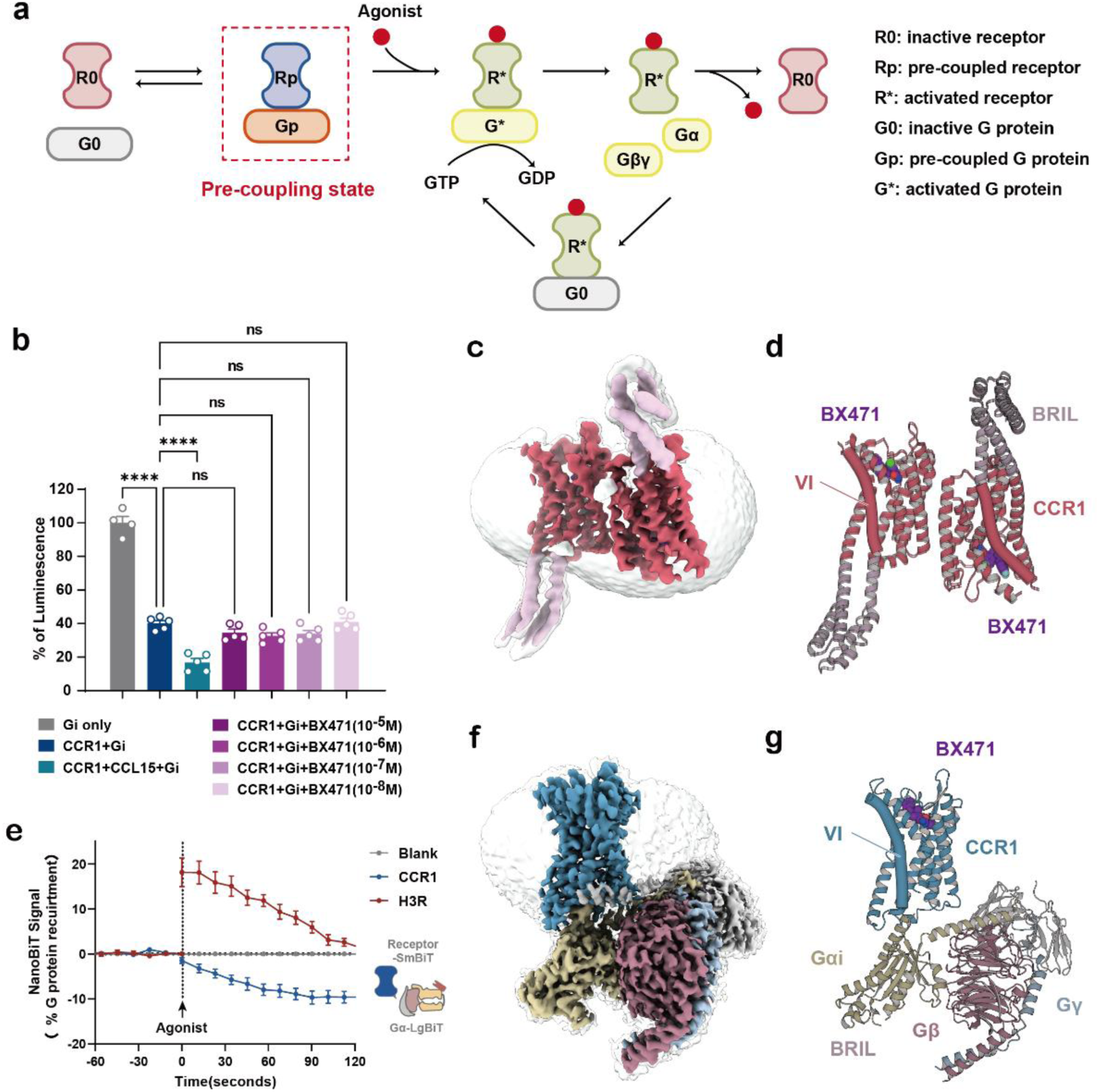
Cryo-EM structures of BX471-CCR1 complexes. **a**, Schematic representation of the GPCR signaling pathway. The canonical GPCR signaling pathway involves several distinct states: the inactive receptor (R0) and G protein (G0), the pre-coupled receptor (Rp) and G protein (Gp), the agonist-bound and the fully active receptor (R*) with the activated G protein (G*). Upon agonist binding, the receptor undergoes conformational changes that promote the exchange of GDP for GTP on the Gα subunit, leading to the dissociation of Gα from the Gβγ dimer and subsequent activation of downstream signaling cascades. **b,** GTP turnover activities of Gi in the presence or absence of CCL15(26-92), BX471, and CCR1. BX471 activities were measured at 10^-5^, 10^-6^, 10^-7^, and 10^-8^ M. The grey column indicates basal activities of Gi proteins alone. Bars and error bars represent the mean and SEM, respectively, of luminescence values corresponding to the residual amounts of GTP after the guanosine triphosphatase reactions. *N* = five independent experiments, performed with single replicates. ns, not significant; ****, P < 0.0001 by one-way ANOVA. **c-d,** Cryo-EM density map (**c**) and model (**d**) of the BX471-CCR1 complex. Rose red, inactive CCR1; light pink, BRIL fusion protein; purple, BX471. **e,** Luciferase complementation between CCR1-SmBit and Gαiβγ-LgBit decreases in response to agonist, whereas luciferase complementation between GPCR (H3R and CCR1)-SmBit and Gαiβγ-LgBit increases in response to agonist. Bars and error bars indicate the mean and SEM. *N* = three independent experiments, performed with single replicates. **f-g,** Cryo-EM density map (**f**) and model (**g**) of the BX471-CCR1-Gi complex. Dodger blue, CCR1 in the SO state; yellow, Gαi1; rosy brown, Gβ; light blue, Gγ; purple, BX471. In **d** and **g**, the TM6 helix is highlighted and shown as a cylinder.

Chemokine receptors, a subfamily of class A GPCRs, are particularly intriguing candidates for studying pre-coupling phenomena^13^. These receptors mediate rapid cellular responses to chemokine gradients, facilitating immune cell migration to sites of inflammation or injury^14,15^. The speed and precision of these responses suggest that pre-coupling might be a common feature among chemokine receptors, potentially contributing to their ability to detect and respond to subtle changes in their chemical environment^16,17^.

In this study, we present the cryo-electron microscopy (cryo-EM) structure of CCR1 in complex with its cognate G protein Gi in a pre-coupled, inactive state. This represents the first structural evidence of a GPCR-G protein complex in the pre-coupled state. Our structure reveals that CCR1 adopts a conformation distinct from both its fully active and inactive states, with the α5 helix of the Gα subunit partially inserted into the receptor’s intracellular cavity. Notably, the C-terminal four amino acids of the Gα subunit are disordered in this pre-coupled state, contrasting with their well-defined α-helical structure in the fully active complex^18–21^. Deletion of the last four residues of the Gα subunit does not affect the formation of the pre-coupled complex but abolishes G protein activation upon agonist binding. This finding indicates that while these residues are not required for G protein recruitment to the receptor, they are essential for the activation process. Thus, the last four amino acids of Gα play a critical role in GPCR-mediated G protein activation but not in the initial pre-coupling interaction.

Furthermore, our results suggest that the ability to form pre-coupled complexes is conserved among chemokine receptors, implying a common mechanism for rapid signal transduction in this GPCR subfamily. The structural insights gained from our study enhance our understanding of the conformational changes that occur during GPCR activation and highlight the specific roles of G protein regions in this process. Overall, our findings provide new molecular insights into the pre-coupling and activation mechanisms of GPCR-G protein interactions. These results have significant implications for the design of drugs targeting specific conformational states of GPCRs and their associated signaling pathways, potentially leading to the development of more selective and efficacious therapeutic agents.

## Results

### Structural determination of CCR1 in the inactive and pre-coupled states

To investigate the structural basis of CCR1 signaling and its interaction with G proteins, we first focused on capturing the receptor in its inactive state. BX471, a potent CCR1 antagonist with Ki values ranging from 1 to 5.5 nM, is widely used in basic research but has not been approved for clinical use (Extended Data Fig. 1a)^22,23^. Our GTP turnover assays revealed that varying concentrations of BX471 did not affect GTP turnover levels (Figure 1b), suggesting that BX471 functions as a neutral antagonist^24,25^. This property makes BX471 an ideal tool for stabilizing CCR1 in its inactive conformation without inducing significant G protein activation.

To determine the structure of inactive CCR1, we employed a strategy using a thermostabilized Escherichia coli apocytochrome b562RIL (BRIL) fused in ICL3 and anti-BRIL Fab (Extended Data Fig. 1b)^26,27^. The cryo-EM structure of BX471-CCR1 in its inactive state was determined at a resolution of 2.9Å, revealing an anti-parallel dimer conformation. Interestingly, despite the use of anti-BRIL Fab during purification, its density was not evident in the final structure, suggesting that stable antibody binding was not crucial for structure determination (Figures 1c, d, and Extended Data Fig. 1c-h).

Building on previous studies showing that some GPCRs form complexes with G proteins before agonist binding, we investigated whether CCR1 could pre-couple with Gi protein. Consistent with earlier fluorescence studies, we observed that stimulation with an endogenous agonist decreased the NanoLuc Binary Technology (NanoBiT) signal between labeled CCR1 and Gi heterotrimers, indicating pre-coupling (Figure 1e)^28^. This observation led us to purify CCR1 in the presence of Gi protein, with BX471 added to stabilize the inactive state (Extended Data Fig. 2a).

Remarkably, we successfully solved the structure of the BX471-CCR1-Gi complex at a resolution of 2.8Å (Figure 2f, g, and Extended Data Fig. 2b-g). This structure revealed several unique features that distinguish it from both the inactive and fully active states of CCR1. In this pre-coupled complex, the receptor stably associated with the G protein without the characteristic outward displacement of TM6 seen in active GPCR-G protein complexes. Notably, the receptor’s helix 8 (H8) was poorly resolved and could not be modeled ab initio. Furthermore, four residues at the C-terminus of Gαi were missing, showing blurred density likely due to substantial flexibility (Extended Data Fig. 2g). These observations align with our functional data showing that these C-terminal residues are critical for G protein activation but not for pre-coupling.

**Figure 2|.**
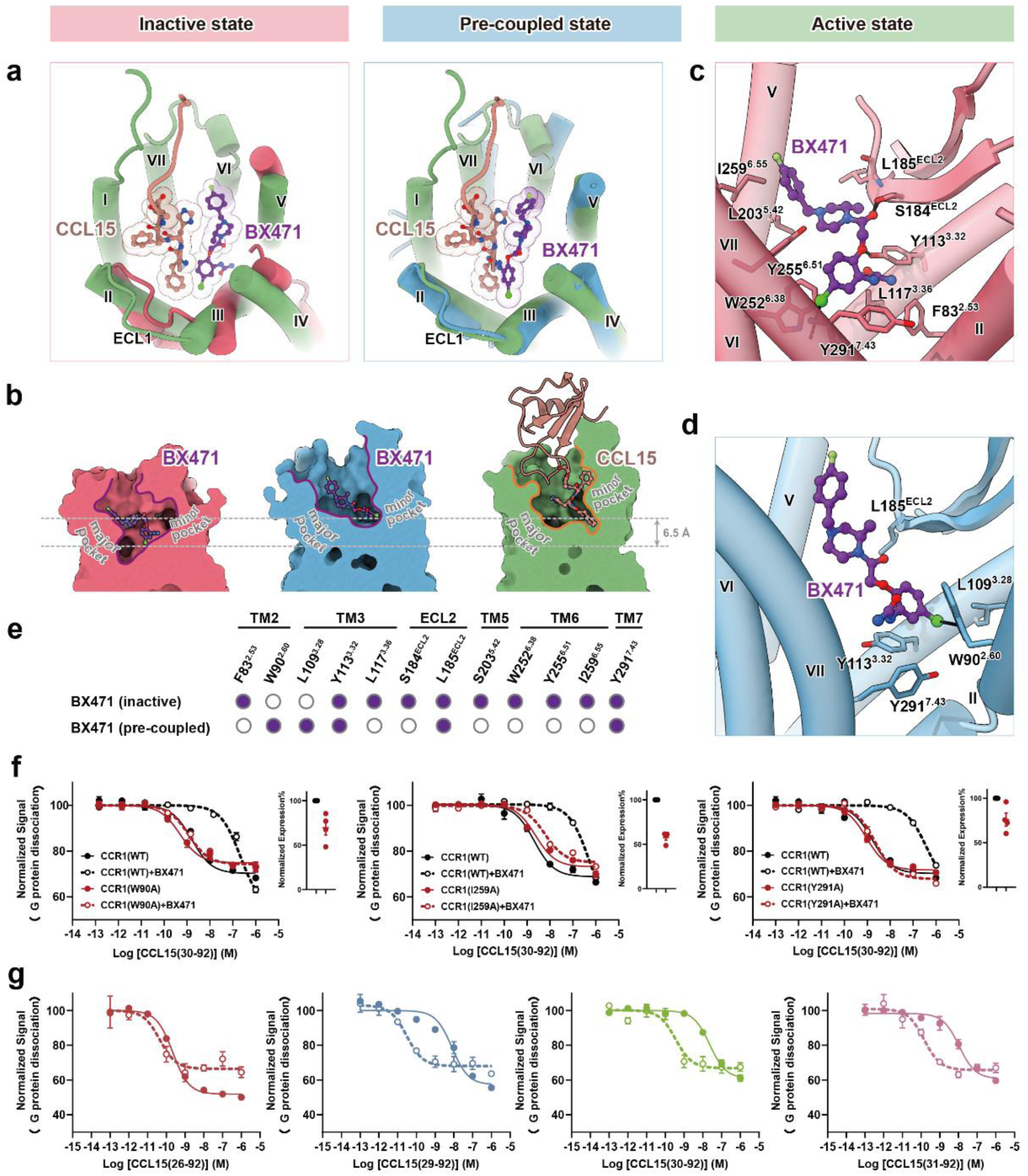
The recognition of BX471 by CCR1. **a**, The orthosteric ligand-binding pocket of BX471-CCR1 (left, rose red) and BX471-CCR1-Gi (right, blue) complexes, aligned with the CCL15(26-92)-CCR1-Gi complex (green, PDB ID: 7VL9). BX471 (purple) and the N-terminal residues of CCL15(26-92) (brown) are shown as spheres. **b,** Surface cut-away views comparing the binding pockets of CCR1 accommodating BX471 (left and middle) and CCL15(26-92) (right). Receptors are shown as surfaces, ligands are shown as diagrams with BX471 and the N-terminus of CCL15(26-92) shown as ball-and-stick models. **c-d,** Detailed interactions between BX471 and CCR1 in the BX471-CCR1 (**c**) and BX471-CCR1-Gi (**d**) complexes. Black dashed lines represent hydrogen bonds. **e,** Residues of CCR1 involved in interactions with BX471 are indicated with solid circles. Residues that show no interaction with the ligand are shown as hollow circles. **f,** Dose-response curves for CCL15(30-92)-induced Gi signaling on wild-type CCR1, CCR1(W90A), CCR1(I259A), and CCR1(Y291A), measured by NanoBiT assay. *N* = 3 independent experiments, performed with duplicate measurements. **g,** NanoBiT G-protein dissociation assays. Dose-response curves for the effects of CCL15 N-terminal variants with or without BX471 (10^-6^ M) treatment. *N* = 3 independent experiments, performed with single replicates. In **f** and **g**, data points and error bars indicate the mean and SEM, respectively.

The pre-coupling of G protein with CCR1 occurs in a manner distinct from the conventional mode observed in all previously known GPCR-G complexes^18–21^. To address concerns that this unique conformation might have resulted from our experimental approach, particularly the use of the LgBiT-HiBiT tagging system to enhance complex stability, we prepared a cryo-EM sample of wild-type CCR1 without this system (Extended Data Fig. 3a)^29,30^. Although the resulting structure was solved at a lower resolution, it confirmed the absence of TM6 outward displacement, validating the authenticity of our observed pre-coupled state (Extended Data Fig.3b-g). Taken together these structural findings provide the first direct evidence for GPCR-G protein pre-coupling and offer new insights into the conformational changes that occur during receptor activation. The unique features of the pre-coupled state, particularly the partial insertion of the G protein’s α5 helix into the receptor’s intracellular cavity and the disorder of the Gα C-terminal residues, suggest a mechanism for rapid signal transduction upon agonist binding.

### The recognition of BX471 by CCR1

Having determined the structures of CCR1 in both inactive and pre-coupled states, we next focused on understanding the molecular basis of BX471 recognition by CCR1. The structures of the BX471-CCR1 complexes revealed the positioning of BX471 within the orthosteric binding pocket of CCR1, which traditionally consists of a minor pocket (formed by TM1-3 and TM7) and a major pocket (formed by TM3-7) (Figure 2a)^31^. Interestingly, unlike the endogenous agonist CCL15(26-92), BX471 occupied both pockets in both the inactive and pre-coupled complexes. However, a notable difference was observed in the binding depth of BX471 between the two states: in the pre-coupled state, BX471 inserted 6.5 Å less deeply into CCR1 compared to the inactive state, although its depth was comparable to that of CCL15 in the active receptor (Figure 2b).

Detailed analysis of the BX471-CCR1 interface in both complexes revealed that the primary interactions centered around BX471’s chloride atom. This observation was consistent with the higher resolution of BX471’s cryo-EM density near the chlorine-terminus compared to the fluorine-terminus (Extended Data Fig. 1h, 2g). In the inactive complex, CCR1 residues Y113^3.32^, S184^ECL^^2^, and Y291^7.43^ formed hydrogen bonds with BX471, while a hydrophobic pocket comprising F83^2.53^, L117^3.36^, L185^ECL2^, L203^5.42^, W252^6.38^, Y255^6.61^, and I259^6.55^ accommodated the ligand (superscript based on Ballesteros-Weinstein numbering rules of GPCRs)^32^. In contrast, the pre-coupled state complex showed a different interaction pattern, with W90^2.60^ of CCR1 forming a hydrogen bond with BX471, and a hydrophobic pocket formed by L109^3.28^, Y113^3.32^, L185^ECL^^2^, and Y291^7.43^ accommodating the ligand (Figure 2c-e). To validate the functional importance of these interactions, we performed alanine substitutions of key binding site residues. Mutations of W90^2.60^A, I259^6.55^A, or Y291^7.43^A significantly reduced BX471’s inhibitory effect on CCR1 activation, supporting our structural observations (Figure 2f, and Extended Data Fig. 4a).

To further explore the pharmacological properties of BX471, we investigated its interactions with various truncated forms of CCL15. Previous studies have shown that under inflammatory conditions, the N-terminus of CCL15 can be proteolytically processed by endogenous enzymes, generating variants with different N-terminal lengths^33,34^. Our experiments revealed a complex pattern of interactions between BX471 and these CCL15 variants. BX471 effectively inhibited CCR1 activation induced by shorter N-terminal variants CCL15(29-92), CCL15(30-92), and CCL15(31-92), but had minimal effect on CCL15(26-92) with its longer N-terminus. Surprisingly, BX471 significantly increased the maximal response (Emax) of CCL15(26-92) by 43.2%. A similar augmentation effect was observed for CCL15(29-92), CCL15(30-92), and CCL15(31-92) at high concentrations (10^-6^ M), where BX471 enhanced their Emax values by 21.9%, 16.8%, and 6.7%, respectively (Figure 2g).

This unexpected enhancement of CCL15-induced responses by BX471 led us to hypothesize that BX471 might be inhibiting CCR1 internalization, thereby increasing cell surface CCR1 levels. To test this hypothesis, we conducted internalization assays, which confirmed that BX471 indeed inhibited CCR1 internalization induced by all tested CCL15 variants (Extended Data Fig. 4b). This finding provides a mechanistic explanation for the observed augmentation effect and highlights the complex pharmacology of CCR1 antagonists. Together, these results collectively demonstrate the intricate nature of ligand recognition by CCR1 and underscore the importance of considering both the binding mode and downstream effects of potential therapeutic compounds. The structural differences in BX471 binding between the inactive and pre-coupled states, coupled with its ability to modulate receptor internalization, offer new insights into the design of CCR1-targeted therapeutics and the potential for biased signaling through this receptor.

### Conformational change of CCR1 in inactive, pre-coupled and active states

To elucidate the activation mechanism of CCR1, we performed a comprehensive comparison of its structures in the inactive, pre-coupled, and active states. This analysis revealed significant conformational changes that provide insights into the molecular basis of CCR1 signaling and the role of pre-coupling in receptor activation.

In the transition from the inactive to the active state, CCR1 underwent substantial displacements in transmembrane helices TM3-7, with the most pronounced changes occurring in TM6. This observation is consistent with the classic GPCR activation model.

Specifically, TM6 rotated outward from the center of the TM bundle to accommodate the α5-helix of Gαi. We quantified this conformational change by measuring the distance between G150^4.42^ and L240^6.46^, which increased from 18.2 Å in the inactive state to 24.4 Å in the active state (Figure 3a-c, and Extended Data Fig. 5a-c).

**Figure 3|.**
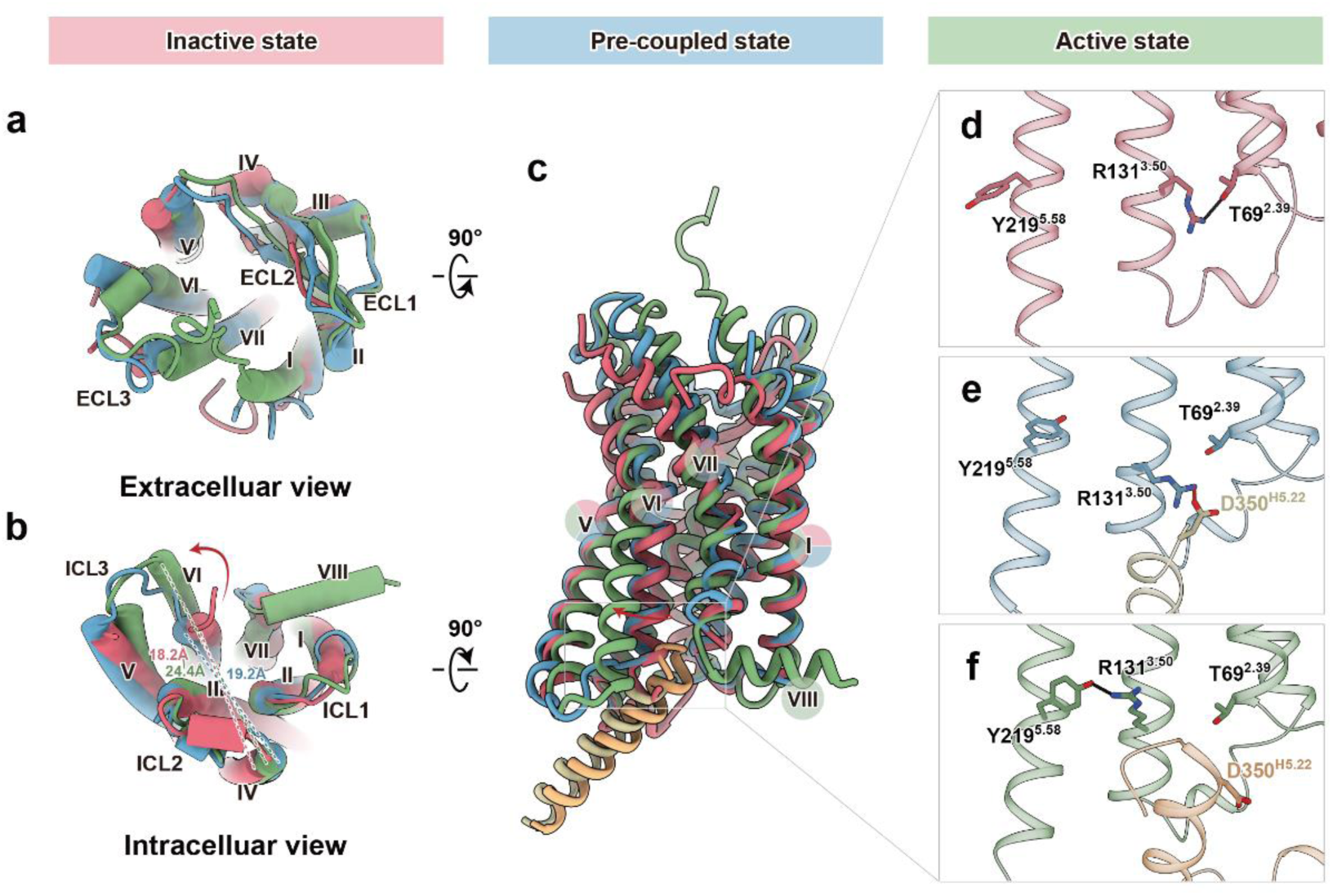
Structural comparison of CCR1 among the inactive, pre-coupled, and active states. **a-c**, Superimposed structures of CCR1 in the pre-coupled state (dodger blue) with inactive CCR1 (rose red) and active CCR1 (green, PDB ID: 7VL9). Extracellular (**a**), intracellular (**b**), and side (**c**) views of the aligned structures are presented. **d-f,** Detailed views of the “ionic lock” region in inactive (**d**), pre-coupled (**e**), and active (**f**) CCR1 structures. Hydrogen bonds are depicted as black dashed lines, while salt bridges are represented by red dashed lines.

Intriguingly, despite stably binding with the G protein, CCR1 in the pre-coupled state adopted a conformation remarkably similar to the inactive state. The G150^4.42^-L240^6.46^ distance in the pre-coupled state measured 19.2 Å, only slightly larger than in the inactive state (Figure 3a-c, and Extended Data Fig. 5d-f). This observation suggests that pre-coupling does not induce the full conformational changes associated with receptor activation. However, subtle differences were noted in the pre-coupled state. The binding pocket for the C-terminal helix of Gαi was formed by conformational changes in the intracellular tips of TM5-7 but remained smaller than in the active state. Consequently, the N-terminal region of CCR1 TM6 (from K233^6.29^ to S235^6.31^) in the pre-coupled state was one helical turn shorter compared to the active state, likely due to steric hindrance with the C-terminus of Gαi. Additionally, the ICL2 region of CCR1 in the inactive state (from A141^ICL^^2^ to A144^ICL^^2^) was one helical turn longer compared to both the active and pre-coupled states (from F140^ICL^^2^ to L142^ICL^^2^), which may contribute to the absence of G protein interaction in the inactive state (Extended Data Fig. 5g-i).

A notable feature of CCR1 is the absence of the conserved “ionic lock” typically found in the inactive state of GPCRs (a salt bridge between R^3.50^ of the DRY motif and D/E^6.30^)^35^. In CCR1, D/E^6.30^ is replaced by lysine. Our structures reveal how this unique feature affects the receptor’s conformational states. In the inactive state, R105^3.50^ formed a hydrogen bond with T69^2.39^ (Figure 3d). In the pre-coupled complex, the G protein-α5 helix partially inserted into CCR1 and formed a salt bridge with R^3.50^, resulting in the opening of the receptor’s cytoplasmic region (Figure 3e). Upon full activation by CCL15, the side chain of R131^3.50^ rotated by approximately 85°, forming hydrogen bonds with Y219^5.58^ of CCR1 (Figure 3f). This series of conformational changes aligns closely with the pre-coupled GPCR-G model proposed by long-scale molecular dynamics (MD) simulations^36^.

To further validate the CCR1 conformation in the pre-coupled state, we performed MD simulations of the receptor with Gα in this state. These simulations provided strong support for the stability of the observed conformation. Throughout the simulations, the CCR1 structure remained globally stable (Extended Data Fig. 6a). While the N-terminus and ECL2 (from Y170^4.62^ to L192^5.31^) showed the highest flexibility, the intracellular portion of TM6 (from K236^6.32^ to L250^6.46^) remained stable (Extended Data Fig. 6b). Importantly, the average distance between G150^4.42^ and L240^6.46^ across all three trajectories was 19.16 ± 0.67 Å, closely matching the value observed in our cryo-EM structure (Figure 3b, and Extended Data Fig. 5f, 6c). These simulations thus provide computational validation for the stability of the unopen TM6 conformation observed in our cryo-EM structure of the pre-coupled state.

These structural and computational analyses collectively provide a detailed picture of the conformational changes CCR1 undergoes during the process of activation. The pre-coupled state emerges as a distinct conformational intermediate, poised for rapid activation upon agonist binding. This mechanism may explain the ability of chemokine receptors to respond swiftly to their ligands, a crucial feature for their function in immune cell migration and inflammatory responses.

### Interaction analysis of the CCR1-G protein interface in pre-coupled and active states

To gain deeper insights into the molecular mechanisms underlying GPCR-G protein coupling, we conducted a detailed analysis of the CCR1-Gi protein interface in both the pre-coupled and active states. This comparison revealed significant differences in the interaction patterns, providing crucial information about the transition from pre-coupling to full activation.

In the pre-coupled state, we observed a distinct orientation of the Gαi α5 helix compared to the active state. The α5 helix was rotated by approximately 14.4° relative to its position in the active CCR1-Gi complex. Notably, the C-terminus of the α5 helix (residues C351^H5.23^ to F354^H5.26^) lacked a distinct density profile in the pre-coupled state (the superscripts following the Gα residues are based on the Common Gα numbering system)^37^ (Figure 4a, and Extended Data Fig. 2g). Low-pass filtered maps further revealed that this C-terminal region was unwound and deflected towards the receptor’s helix 8 (H8), with density too weak to support model building (Extended Data Fig. 7a-b). These observations suggest that while the C-terminus of the Gαi helix contributes to the formation of the GPCR-G complex, it is not essential for pre-coupling. This finding aligns with previous studies on β_2_AR-mediated Gs activation, where time-resolved structural mass spectrometry showed that the C-terminus of the α5 helix remained dynamic during receptor-G protein coupling^5^. Interestingly, despite these differences, the overall structures of Gαi in both active and pre-coupled complexes were similar. In both states, the α-helical domain of Gαi adopted an open conformation distinct from the GDP-bound state, indicating a nucleotide-free state in both the active and pre-coupled complexes (Extended Data Fig. 7c-d).

**Figure 4|.**
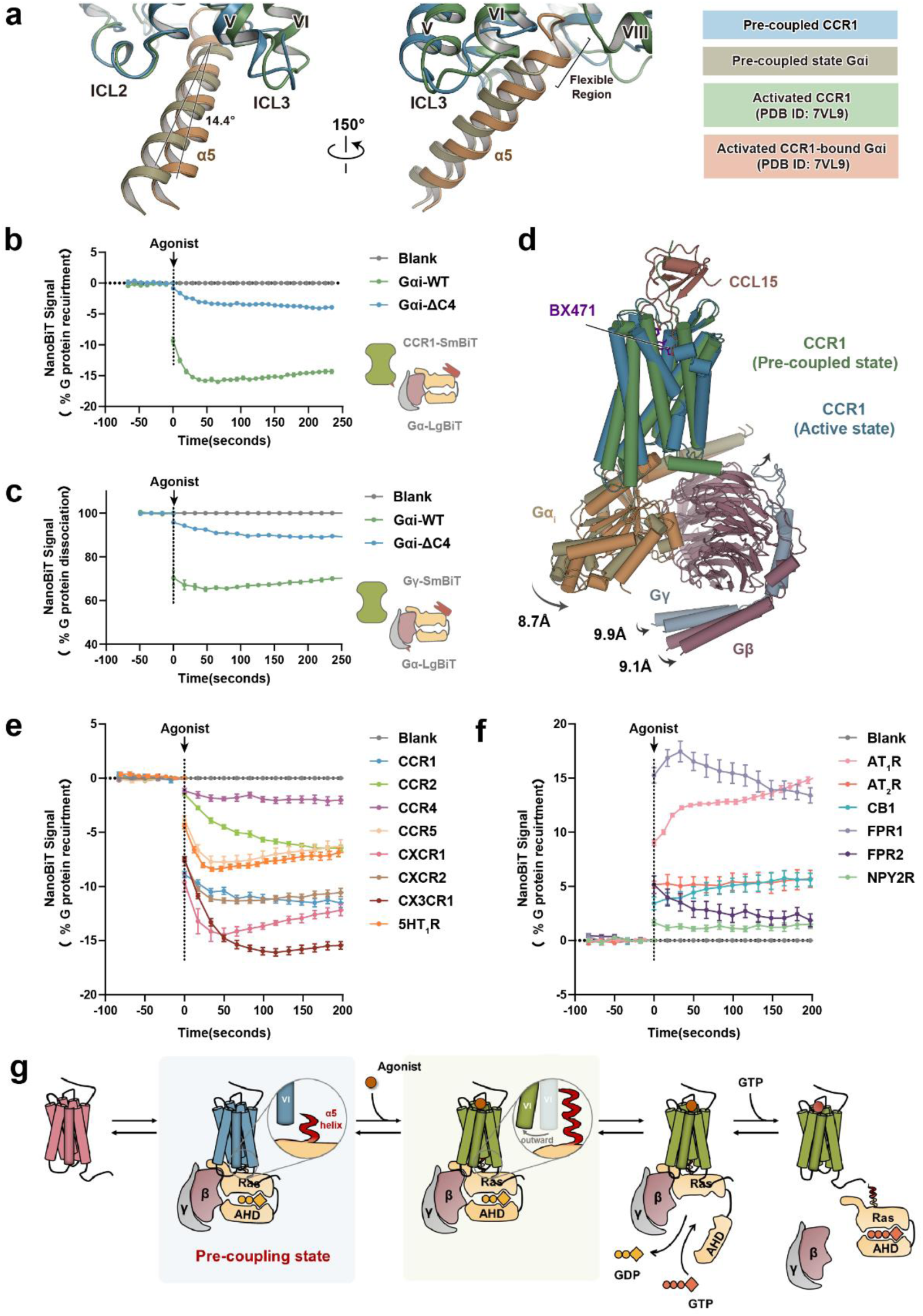
Comparison of CCR1-G protein interfaces in active and pre-coupled complexes. **a**, Comparison of the interface between the α5 helix of Gαi and the cytoplasmic region of CCR1 in the active (green) and pre-coupled (dodger blue) states. The alignment is performed using the receptor as the reference. **b-c,** Agonist-induced changes in luciferase signal measured by NanoBiT between CCR1-SmBiT and Gα-LgBiT (**b**), and between Gγ-SmBiT and Gα-LgBiT (**c**). Bars and error bars indicate the mean and SEM, respectively. *N* = 6 (**b**) and 4 (**c**) independent experiments, performed with single replicates. **d,** Orthogonal views of the structures of the CCR1-Gi heterotrimer complex in the active and pre-coupled states, colored by subunit. CCR1 is shown in green (active state) and dodger blue (pre-coupled state), Gαi1 in gold (active state) and yellow (pre-coupled state), Gβ in rosy brown, Gγ in light blue, and CCL15 in brown. **e-f,** Luciferase complementation between various GPCRs (CCR1, CCR2, CCR4, CCR5, CXCR1, CXCR2, CX3CR1, and 5HT_1_R) fused to SmBiT and Gα-LgBiT decreases in response to agonist (**e**), whereas luciferase complementation between other GPCRs (NPY2R, FPR1, FPR2, CB1, AT_1_R, and AT_2_R) fused to SmBiT and Gα-LgBiT increases in response to agonist (**f**). *N* = 6-8 independent experiments, performed with single replicates. **g,** The model of GPCR activation in which the C-terminal residues of the Gαi α5 helix (red), while disordered in the pre-coupled state (blue), play a crucial role in the transition to the fully active state (green) and subsequent G protein dissociation

To further investigate the functional role of the Gαi C-terminus, we employed the NanoBiT system to measure agonist-induced dissociation of CCR1 from Gαi. Agonist stimulation led to a decrease in NanoBiT signal between CCR1-SmBiT and LgBiT-tagged wild-type Gαi, indicating dissociation. However, truncation of the last 4 amino acids of Gαi (Gαi-ΔC4) abolished this dissociation (Figure 4b). Similar results were observed for the dissociation of Gβγ from Gαi (Figure 4c). These findings demonstrate that while the C-terminus of Gα is not essential for pre-coupling, it plays a crucial role in G protein dissociation and activation of downstream signaling pathways upon agonist binding.

Structural superimposition of active and pre-coupled complexes revealed a similar interface within the ICL2 region (Extended Data Fig. 8a). Both complexes showed direct interactions between numerous critical sites at ICL2 and the C-terminus of TM3 with Gαi, including a hydrogen bond at A134^3.53^ and hydrophobic interactions involving I135^3.54^, A138^ICL2^, and L142^ICL2^ (Extended Data Fig. 8b, c). However, the G-protein interface with ICL3 and the intracellular ends of TM5 and TM6 differed between the active and pre-coupled states (Extended Data Fig. 8a). The pre-coupled state complex featured an additional hydrogen bond between L348^H5.20^ and A237^6.33^, potentially compensating for the absence of interactions with H8 (Extended Data Fig. 8d, e). These differences resulted in an approximately 9Å deflection of the Gαβγ heterotrimer in the pre-coupled CCR1-Gi complex compared to the active state, highlighting the plasticity in G-protein binding modes (Figure 4d).

Sequence alignment of class A GPCRs revealed that the residues involved in CCR1 pre-coupling with Gi are conserved among many chemokine receptors, including CCR2, CCR4, CCR5, CXCR1, CXCR2, and CX3CR1 (Extended Data Fig. 9a-b). Agonist stimulation decreased the NanoBiT signal for these receptors, confirming that these receptors could pre-couple with G protein prior to agonist binding. Interestingly, 5-HT_1_R, which shares high sequence similarity with CCR1, also demonstrated pre-coupling capacity (Figure 4e). In contrast, other Gi-coupled GPCRs with low sequence similarity to CCR1, such as NPY2R, FPR1, FPR2, CB1, AT_1_R, and AT_2_R, recruited G protein only after agonist binding. This phenomenon suggests no pre-coupling (Figure 4f, and Extended Data Fig. 9c-d).

These comprehensive analyses of the CCR1-Gi interface and the broader examination of pre-coupling across GPCR families underscore the widespread nature of this phenomenon, particularly among chemokine receptors. The structural and functional insights gained from this study provide a foundation for understanding the molecular basis of rapid GPCR signaling and open new avenues for drug discovery targeting specific conformational states of GPCR-G protein complexes.

## Discussion

Our study provides the first structural evidence of GPCR-G protein pre-coupling, offering crucial insights into the molecular mechanisms underlying rapid signal transduction in chemokine receptors. By capturing the structure of the CCR1-Gi complex in its pre-coupled state, we have revealed a distinct conformational intermediate that bridges our understanding of GPCR activation from the inactive to the fully active state.

The pre-coupled state of CCR1-Gi demonstrates several unique features that distinguish it from both the inactive and fully active conformations. Most notably, the α5 helix of Gαi partially penetrates the intracellular region of CCR1, forming a salt bridge with R^3.50^ of the conserved DRY motif. This interaction disrupts the coupling between transmembrane segments and partially opens the cytoplasmic region of the receptor, priming it for rapid activation upon agonist binding. Interestingly, the C-terminal residues of the Gαi α5 helix, while disordered in the pre-coupled state, play a crucial role in the transition to the fully active state and subsequent G protein dissociation, as evidenced by our functional studies with the Gαi-ΔC4 mutant.

The conformational changes observed in CCR1 during pre-coupling are subtle yet significant. The distance between G150^4.42^ and L240^6.46^, a key measure of TM6 movement, increases only slightly in the pre-coupled state compared to the inactive state. This suggests that pre-coupling prepares the receptor for activation without fully engaging the conformational changes associated with agonist binding. The stability of this pre-coupled conformation, supported by our molecular dynamics simulations, indicates that it represents a distinct, physiologically relevant state in the GPCR activation process.

Our structural analysis also revealed important differences in the CCR1-Gi interface between the pre-coupled and active states. While both states share similar interactions in the ICL2 region, the pre-coupled state exhibits unique interactions around ICL3 and the intracellular ends of TM5 and TM6. These differences result in a significant deflection of the Gαβγ heterotrimer in the pre-coupled complex, highlighting the plasticity of G protein binding modes and suggesting a mechanism for the transition to the fully active state upon agonist binding.

The conservation of residues involved in pre-coupling among chemokine receptors, as revealed by our sequence alignment and functional studies, suggests that this phenomenon may be a common feature of this GPCR subfamily. This conservation could explain the ability of chemokine receptors to respond rapidly to their ligands, a crucial aspect of their function in immune cell migration and inflammatory responses. The observation that other Gi-coupled receptors with low sequence similarity to CCR1 do not exhibit pre-coupling further underscores the specificity of this mechanism to chemokine receptors and closely related GPCRs like 5-HT_1_R.

Our findings also have important implications for drug discovery. The structure of the pre-coupled CCR1-Gi complex reveals a unique conformational state that could serve as a novel target for therapeutic interventions. By designing compounds that specifically stabilize or destabilize the pre-coupled state, it may be possible to modulate receptor activity with greater precision. This approach could be particularly valuable for developing treatments for inflammatory and autoimmune conditions where chemokine receptor signaling plays a crucial role.

The study of BX471 binding to CCR1 in both the inactive and pre-coupled states provides additional insights into ligand recognition and receptor pharmacology. The differential binding mode of BX471 in these two states, coupled with its ability to modulate receptor internalization, highlights the complex nature of GPCR-ligand interactions. These observations suggest that the development of biased ligands targeting specific conformational states of chemokine receptors may be a promising avenue for therapeutic development.

In conclusion, our study presents a comprehensive model of GPCR activation that incorporates the pre-coupled state as a key intermediate. This model suggests that GPCR coupling with G proteins can occur before agonist-induced activation, with the pre-coupled complex remaining in a primed yet resting state until agonist binding triggers full activation. This mechanism provides a molecular basis for the rapid and sensitive responses characteristic of chemokine receptor signaling (Figure 4g).

These findings not only advance our fundamental understanding of GPCR biology but also open new avenues for drug discovery. By targeting specific conformational states of GPCR-G protein complexes, it may be possible to develop more selective and efficacious therapeutics for a range of diseases involving chemokine signaling. Future studies exploring the prevalence and functional significance of pre-coupling across other GPCR families will undoubtedly yield further insights into this important aspect of cellular signal transduction.

## Methods

### Expression and Purification of 3C and TEV Protease

The 3C and TEV protease expression strains (kindly provided by Prof. Yan Zhang’s group at Zhejiang University) were amplified separately in 4 L of LB liquid medium containing ampicillin (100 μg/mL) and chloramphenicol (34 μg/mL). The cultures were incubated at 37°C with shaking at 220 rpm until the OD600 reached 0.6-0.8. Protein expression was induced by adding isopropyl-β-D-thiogalactopyranoside (IPTG) to a final concentration of 0.3 mM, followed by incubation at 18°C with shaking for 20 hours. The bacterial cells were harvested by centrifugation at 3,000g for 15 minutes, and the pellet was resuspended in 200 mL of lysis buffer (20 mM HEPES pH 8.0, 300 mM NaCl, 2 mM MgCl2) supplemented with lysozyme (0.5 mg/mL) and phenylmethylsulfonyl fluoride (PMSF, 1 mM). The cells were lysed using a high-pressure homogenizer, and the lysate was clarified by centrifugation at 30,000g for 30 minutes at 4°C. The supernatant was collected, and imidazole was added to a final concentration of 30 mM. The clarified lysate was loaded onto a 5 mL nickel-nitrilotriacetic acid (Ni-NTA) resin (GE Healthcare) pre-equilibrated with lysis buffer. The column was washed with 50 mL of wash buffer (20 mM HEPES pH 8.0, 300 mM NaCl, 2 mM MgCl2, 30 mM imidazole) to remove non-specifically bound proteins. The target protein was eluted with an appropriate volume of elution buffer (20 mM HEPES pH 8.0, 300 mM NaCl, 2 mM MgCl2, 250 mM imidazole), and the elution fractions were collected. The eluted protein was concentrated to approximately 500 μL using a 30 kDa molecular weight cut-off centrifugal filter unit (Millipore). The concentrated protein was further purified by size exclusion chromatography using a Superdex 200 16/60 column (GE Healthcare) equilibrated with SEC buffer (20 mM HEPES pH 8.0, 100 mM NaCl). The fractions containing the purified proteases were pooled and concentrated to 10 mg/mL for 3C protease and 1.5 mg/mL for TEV protease. Glycerol was added to a final concentration of 10% (w/v) as a cryoprotectant. The purified proteins were aliquoted, flash-frozen in liquid nitrogen, and stored at -80°C until further use.

### Expression and Purification of CCL15 N-Terminal Variants

To explore the structure-function relationship of CCL15, we engineered several N-terminal truncation variants, ranging from CCL15(26-92) to CCL15(31-92). Each variant was designed to include a GP64 signal peptide for secretion and a C-terminal maltose-binding protein (MBP) fused with an 8×His-tag to facilitate expression and purification. The variants were expressed using the bac-to-bac expression system in Sf9 insect cells (Invitrogen), and purified through Ni-NTA affinity chromatography and size exclusion chromatography (SEC). The expression of the CCL15(26-92)-MBP construct was carried out in Sf9 cells cultured in protein-free ESF 921 insect cell culture medium (Expression Systems). The cells were infected with the recombinant baculovirus at a multiplicity of infection (MOI) of 100 and incubated for 48 hours at 27°C. The cell culture supernatant was collected and loaded onto a Ni-NTA resin (GE Healthcare) pre-equilibrated with binding buffer (20 mM Tris-HCl, 500 mM NaCl, 5 mM imidazole, pH 7.5). The column was washed with 20 column volumes of wash buffer (20 mM Tris-HCl, 500 mM NaCl, 20 mM imidazole, pH 7.5) and eluted with elution buffer (20 mM Tris-HCl, 500 mM NaCl, 500 mM imidazole, pH 7.5). The eluted protein was then treated with 3C protease at a ratio of 1:100 (w/w) in the presence of 10% glycerol to cleave the MBP tag. After incubating the mixture for 12 hours at 4°C, the resulting protein was concentrated to a final volume of 500 μL using an Amicon Ultra-15 Centrifugal Filter Unit with a 3 kDa molecular weight cut-off (Millipore). The protein was further purified through SEC using a Superdex™ 75 Increase 10/300 GL column (GE Healthcare) in a buffer containing 20 mM HEPES (pH 7.5), 100 mM NaCl, and 10% glycerol. The column was equilibrated and run at a flow rate of 0.5 mL/min. Monomeric fractions were identified based on the elution volume and SDS-PAGE analysis, collected, and concentrated using the aforementioned centrifugal filter unit. Protein concentration was determined using a NanoDrop™ 2000 spectrophotometer (Thermo Scientific) with the protein’s theoretical extinction coefficient. The purified proteins were flash-frozen in liquid nitrogen and stored at -80°C until biochemical assays. The same procedure was applied to produce other CCL15 variants, from CCL15(27-92) to CCL15(31-92).

### Expression and Purification of CCR1

The wild-type human CCR1 cDNA was cloned into a modified pFastBac1 vector (Invitrogen), incorporating an N-terminal FLAG tag and a C-terminal 8×His-tag for affinity purification. The recombinant CCR1 baculovirus was generated using the Bac-to-Bac® Baculovirus Expression System (Invitrogen). For protein expression, Sf9 insect cells (Invitrogen) were grown in protein-free ESF 921 insect cell culture medium (Expression Systems) to a density of 2.8×10^6 cells/mL in a 2 L shaker flask (Corning). The cells were then infected with the recombinant CCR1 baculovirus at a MOI of 100. After 48 hours of incubation at 27°C with shaking at 130 rpm, the cells were harvested by centrifugation at 1,000g for 15 minutes at 4°C. The cell pellets were flash-frozen in liquid nitrogen and stored at −80°C until protein extraction. To extract the CCR1 protein, the frozen cell pellets were thawed and resuspended in lysis buffer containing 20 mM HEPES (pH 7.5), 2 mM MgCl2, 100 mM NaCl, and a protease inhibitor cocktail (Roche, Complete™ EDTA-free). Membranes were solubilized by adding lauryl maltose neopentyl glycol (LMNG, Anatrace) and cholesteryl hemisuccinate (CHS, Anatrace) to final concentrations of 0.5% (w/v) and 0.1% (w/v), respectively. The mixture was incubated for 2 hours at 4°C with gentle agitation. Insoluble debris was removed by centrifugation at 30,000g for 30 minutes at 4°C. The supernatant containing solubilized CCR1 was supplemented with imidazole (Sigma-Aldrich) to a final concentration of 20 mM and then loaded onto a Ni-NTA resin (GE Healthcare) pre-equilibrated with binding buffer (20 mM HEPES pH 7.5, 100 mM NaCl, 20 mM imidazole, 0.01% LMNG, and 0.002% CHS). The column was washed with 20 column volumes of binding buffer, and the bound proteins were eluted with elution buffer (20 mM HEPES pH 7.5, 100 mM NaCl, 500 mM imidazole, 0.01% LMNG, and 0.002% CHS). The Ni-NTA-purified CCR1 was further purified by anti-FLAG M1 affinity chromatography. The Ni-NTA elution fractions were pooled, supplemented with CaCl2 to a final concentration of 3 mM, and then incubated with anti-FLAG M1 affinity resin (Sigma-Aldrich) for 2 hours at 4°C. The resin was washed with 10 column volumes of washing buffer (20 mM HEPES pH 7.5, 100 mM NaCl, 2 mM MgCl2, 3 mM CaCl2, 0.01% LMNG, and 0.002% CHS). The bound CCR1 was eluted with elution buffer (20 mM HEPES pH 7.5, 100 mM NaCl, 2 mM MgCl2, 0.01% LMNG, 0.002% CHS, 0.1 mg/ml FLAG peptide, and 5 mM EGTA). The eluted CCR1 fractions were concentrated using a 50 kDa molecular weight cut-off Amicon Ultra centrifugal filter unit (Millipore). The concentrated protein was flash-frozen in liquid nitrogen and stored at −80°C until further use.

### Expression and Purification of Gi1 Heterotrimer

The human Gαi1 subunit was cloned into a pFastBac1 vector (Invitrogen), while the N-terminal 6×His-tagged wild-type human Gβ1 and untagged Gγ2 subunits were cloned into a pFastBac-Dual vector (Invitrogen). Recombinant baculoviruses were generated using the Bac-to-Bac® Baculovirus Expression System (Invitrogen). For protein expression, Hi5 insect cells (Invitrogen) were grown in ESF 921 insect cell culture medium (Expression Systems) supplemented with L-glutamine to a density of 3.0×10^6 cells/ml in a 2 L shaker flask (Corning). The cells were then co-infected with the recombinant Gαi1 and Gβ1γ2 baculoviruses at a volumetric ratio of 1:1 and a total MOI of 100. After incubation for 48 hours at 27°C with shaking at 120 rpm, the cells were harvested by centrifugation at 1,000g for 15 minutes at 4°C. The cell pellets were flash-frozen in liquid nitrogen and stored at −80°C until purification. For Gi1 heterotrimer purification, the frozen Hi5 cell pellets were thawed and resuspended in lysis buffer (10 mM HEPES pH 7.5, 100 μM MgCl2, 10 μM GDP, and cOmplete™ Protease Inhibitor Cocktail (Roche)). The cell membranes were isolated by homogenization using a Dounce homogenizer followed by centrifugation at 150,000g for 30 minutes at 4°C. The isolated membranes were solubilized in solubilization buffer (1% sodium cholate, 0.05% DDM, 10 μM GDP, 2 mM MgCl2, 20 mM HEPES pH 7.5, and 100 mM NaCl) for 1 hour at 4°C with gentle agitation. The solubilized membrane fraction was collected by centrifugation at 150,000g for 30 minutes at 4°C. The supernatant containing the solubilized Gi1 heterotrimer was incubated with a Ni-NTA resin (GE Healthcare) pre-equilibrated with solubilization buffer. The resin was then washed with 20 column volumes of wash buffer (20 mM HEPES pH 7.5, 100 mM NaCl, 0.1% LMNG, 20 mM imidazole, 10 μM GDP, and 2 mM MgCl2). During the washing steps, the detergent was exchanged from 1% sodium cholate and 0.05% DDM to 0.1% LMNG to stabilize the heterotrimer. The bound Gi1 heterotrimer was eluted with elution buffer (20 mM HEPES pH 7.5, 100 mM NaCl, 0.1% LMNG, 250 mM imidazole, 10 μM GDP, and 2 mM MgCl2). The eluted Gi1 heterotrimer was then incubated with His-tagged TEV protease at a 1:10 (w/w) ratio overnight at 4°C to remove the N-terminal 6×His-tag from the Gβ1 subunit. The cleaved heterotrimer was then passed through a Ni-NTA resin to remove the cleaved His-tag and the His-tagged TEV protease. The purified Gi1 heterotrimer was concentrated using a 50 kDa molecular weight cut-off Amicon Ultra centrifugal filter unit (Millipore) and further purified by size-exclusion chromatography using a Superose™ 6 Increase 10/300 GL column (Cytiva) equilibrated with running buffer (20 mM HEPES pH 7.5, 100 mM NaCl, 0.01% LMNG, 2 mM MgCl2, and 10 μM GDP). The fractions containing the purified Gi1 heterotrimer were concentrated, flash-frozen in liquid nitrogen, and stored at −80°C until use.

### GTPase-Glo Assay

The GTPase activity of CCR1-coupled Gi1 heterotrimer was measured using the GTPase-Glo™ Assay kit (Promega). Purified CCR1 and Gi1 heterotrimer were prepared as described in the previous sections. For the assay, three different conditions were tested: unliganded CCR1, CCL15-bound CCR1, and BX471-bound CCR1. The small-molecule antagonist BX471 was obtained from MedChemExpress (Cat. No. HY-12080). Purified CCR1 (4 µM final concentration) was pre-incubated with either 1 µM CCL15(26-92), BX471 (final concentrations of 10, 1, 0.1, and 0.01 µM), or buffer alone (20 mM HEPES pH 7.5, 100 mM NaCl, 0.02% LMNG) for 30 minutes at room temperature. The GTPase reaction was initiated by mixing the pre-incubated CCR1 with purified Gi1 heterotrimer (5 µM final concentration) in a 384-well white opaque plate (Corning) in a total volume of 5 µL. The final reaction buffer contained 20 mM HEPES pH 7.5, 100 mM NaCl, 0.02% LMNG, 1 mM MgCl2, 5 µM GTP, and 5 µM GDP.

The GTPase reaction mixture was incubated at room temperature for 2 hours. After incubation, 5 µL of reconstituted GTPase-Glo™ Reagent (1X) was added to each well using a multichannel pipette. The plate was mixed briefly by orbital shaking at 500 rpm for 30 seconds and then incubated at room temperature for 30 minutes to convert the remaining GTP to ATP. Following the 30-minute incubation, 10 µL of Detection Reagent was added to each well using a multichannel pipette. The plate was mixed briefly by orbital shaking at 500 rpm for 30 seconds and then incubated at room temperature for 5-10 minutes to allow for luminescence signal stabilization. The luminescence signal was measured using a TECAN multimode microplate reader with an integration time of 500 ns per well. The raw luminescence data were analyzed using GraphPad Prism 9 software. Background luminescence (from a control reaction without Gi1 heterotrimer) was subtracted from all sample readings, and the data were normalized to the unliganded CCR1 condition.

### Purification of scFv16 and Anti-thermostabilized apocytochrome b562RIL (BRIL) Fab

The scFv16 construct was engineered to include a GP64 signal peptide for secretion and a C-terminal 8×His-tag for purification (kindly provided by Prof. Yan Zhang’s group at Zhejiang University). The protein was expressed using the bac-to-bac expression system in Hi5 insect cells (Invitrogen). The cells were infected with the recombinant baculovirus at a multiplicity of infection (MOI) of 100 and incubated for 48 hours at 27°C. The cell culture supernatant containing the secreted scFv16 was collected and loaded onto a Ni-NTA resin (GE Healthcare) pre-equilibrated with binding buffer (20 mM HEPES pH 7.5, 100 mM NaCl, 20 mM imidazole). The column was washed sequentially with high-salt Ni wash buffer (20 mM HEPES pH 7.5, 500 mM NaCl, 20 mM imidazole) and low-salt Ni wash buffer (20 mM HEPES pH 7.5, 100 mM NaCl, 20 mM imidazole). The bound protein was eluted with Ni elution buffer (20 mM HEPES pH 7.5, 100 mM NaCl, 250 mM imidazole). The eluted scFv16 protein was further purified by size exclusion chromatography using a Superdex 200 Increase 10/300 GL column (GE Healthcare) equilibrated with SEC buffer (20 mM HEPES pH 7.5, 100 mM NaCl). Monomeric fractions were collected, concentrated using a 10 kDa molecular weight cut-off centrifugal filter unit (Millipore), flash-frozen in liquid nitrogen, and stored at -80°C until further use.

For anti-BRIL Fab purification, the light chain (LC) and heavy chain (HC) sequences were cloned into separate pFastBac1 vectors, each containing a GP64 signal peptide for secretion (kindly provided by Prof. H. Eric Xu’s group at Chinese Academy of Sciences). An 8×His-tag was added to the C-terminus of the LC for purification purposes. High Five insect cells were co-infected with equal MOIs of recombinant baculoviruses containing anti-BRIL Fab HC and LC. The cells were incubated for 48 hours at 27°C, and the cell culture supernatant was collected.The anti-BRIL Fab was purified from the supernatant using the same protocol as described for scFv16. Briefly, the supernatant was loaded onto a Ni-NTA resin, washed with high-salt and low-salt Ni wash buffers, and eluted with Ni elution buffer. The eluted anti-BRIL Fab was further purified by size exclusion chromatography using a Superdex 200 Increase 10/300 GL column equilibrated with SEC buffer (20 mM HEPES pH 7.5, 100 mM NaCl). Monomeric fractions were collected, concentrated using a 30 kDa molecular weight cut-off centrifugal filter unit (Millipore), flash-frozen in liquid nitrogen, and stored at -80°C until further use.

### Expression and Purification of BX471-CCR1-Gi Complex

The BX471-CCR1-Gi complex was purified using two different approaches, depending on the inclusion of the HiBiT system. Initially, the apo CCR1-Gi complex was purified as previously described^29^. For the version of the BX471-CCR1-Gi complex without the HiBiT system, an additional TEV protease cleavage site was introduced between the receptor and the LgBiT subunit, allowing the removal of the HiBiT system through TEV treatment. The TEV protease cleavage site between the receptor and the LgBiT subunit enabled the removal of the HiBiT components, yielding a more native-like BX471-CCR1-Gi complex. After completing size exclusion chromatography, BX471 (MedChemExpress, Cat. No. HY-12080) was dissolved in a buffer containing 20 mM HEPES pH 7.5, 100 mM NaCl, 0.0075% (w/v) lauryl maltose neopentyl glycol (LMNG, Anatrace), 0.002% (w/v) cholesteryl hemisuccinate (CHS, Anatrace), and 0.0025% (w/v) glyco-diosgenin (GDN, Anatrace) to a final concentration of 100 μM. A 600 μL aliquot of the apo CCR1-Gi complex fraction was combined with 45 μL of the BX471 solution and incubated for 30 minutes at 4°C to facilitate complex formation. Following incubation, the sample was concentrated to a volume of 50 μL using a 100 kDa molecular weight cut-off centrifugal filter unit (Millipore). The concentrated sample was then diluted 10-fold with a buffer containing 20 mM HEPES pH 7.5 and 100 mM NaCl to reduce the detergent concentration. The diluted sample was concentrated once more using the same centrifugal filter unit to a final volume suitable for cryo-electron microscopy grid preparation.

### Expression and Purification of Inactive BX471-CCR1 Complex

To purify the inactive BX471-CCR1 complex, the intracellular loop 3 (ICL3) of CCR1 (residues Arg229-Asn231) was replaced with a BRIL. The resulting CCR1-BRIL construct, which included an N-terminal hemagglutinin (HA) signal sequence, a FLAG epitope tag, and a C-terminal 10×His tag, was cloned into a pFastBac1 vector (Invitrogen). Baculovirus production and protein expression were performed using the bac-to-bac expression system. Sf9 insect cell cultures (Expression Systems) were grown in ESF 921 insect cell culture medium (Expression Systems) to a density of 2.7×10^6 cells/mL and then infected with the recombinant baculovirus encoding the CCR1-BRIL construct at a multiplicity of infection (MOI) of 100. The infected cells were incubated at 27°C for 48 hours, harvested by centrifugation at 1,000g for 15 minutes, and stored at -80°C until further use. BX471 (MedChemExpress, Cat. No. HY-12080) was prepared in a buffer containing 20 mM HEPES pH 7.5, 100 mM NaCl, 0.0075% (w/v) LMNG (Anatrace), 0.002% (w/v) CHS (Anatrace), and 0.0025% (w/v) GDN (Anatrace) to a final concentration of 100 μM. Frozen insect cell pellets were thawed and resuspended in lysis buffer (20 mM HEPES pH 7.5, 2 mM MgCl2, 100 mM NaCl, EDTA-free Protease Inhibitor Cocktail (Biomake), anti-BRIL Fab, and 1 μM BX471). Membranes were solubilized by adding LMNG and CHS to final concentrations of 0.5% (w/v) and 0.1% (w/v), respectively, followed by incubation for 3 hours at 4°C with gentle agitation. The solubilized membranes were clarified by centrifugation at 30,000g for 30 minutes at 4°C. The supernatant was then incubated with Ni-NTA resin (GE Healthcare) pre-equilibrated with lysis buffer for 2 hours at 4°C. The resin was washed with 20 column volumes of Ni wash buffer (20 mM HEPES pH 7.5, 100 mM NaCl, 2 mM MgCl2, 0.01% (w/v) LMNG, 0.002% (w/v) CHS, 1 μM BX471, and 30 mM imidazole). The bound protein was eluted with Ni elution buffer (20 mM HEPES pH 7.5, 100 mM NaCl, 2 mM MgCl2, 0.01% (w/v) LMNG, 0.002% (w/v) CHS, 1 μM BX471, and 250 mM imidazole). The Ni-NTA eluate was supplemented with 3 mM CaCl2 and applied to M1 anti-FLAG immunoaffinity resin (Sigma-Aldrich) pre-equilibrated with FLAG wash buffer (20 mM HEPES pH 7.5, 100 mM NaCl, 2 mM MgCl2, 0.01% (w/v) LMNG, 0.002% (w/v) CHS, 1 μM BX471, and 3 mM CaCl2). The resin was washed with 10 column volumes of FLAG wash buffer, and the bound complex was eluted with FLAG elution buffer (20 mM HEPES pH 7.5, 100 mM NaCl, 2 mM MgCl2, 0.01% (w/v) LMNG, 0.002% (w/v) CHS, 1 μM BX471, and 0.2 mg/mL 3×FLAG peptide (Beyotime, Cat. No. P9801)). The FLAG eluate was further purified by size exclusion chromatography using a Superose 6 Increase 10/300 GL column (GE Healthcare) equilibrated with SEC buffer (20 mM HEPES pH 7.5, 100 mM NaCl, 0.00075% (w/v) LMNG, 0.0002% (w/v) CHS, 0.00025% (w/v) GDN, and 1 μM BX471). Monomeric fractions were pooled, concentrated using a 100 kDa molecular weight cut-off centrifugal filter unit (Millipore), and prepared for cryo-electron microscopy grid preparation.

### Cryo-EM grid preparation and data collection

For cryo-EM grid preparation, 3 μL of purified BX471-CCR1 or BX471-CCR1-Gi complex at a concentration of approximately 15 mg/mL were applied onto glow-discharged holey carbon grids (Quantifoil, R1.2/1.3). The grids were blotted for 3.5 seconds with a blot force of -2 at 100% humidity and 4°C, then plunge-frozen in liquid ethane using a Vitrobot Mark IV (Thermo Fisher Scientific). The frozen grids were transferred to liquid nitrogen for storage until data collection.

Cryo-EM data for the BX471-CCR1-Gi complex with the HiBiT system were collected at the Center of Cryo-Electron Microscopy, Zhejiang University (Hangzhou, China). Cryo-EM data for the complexes of inactive BX471-CCR1 and BX471-CCR1-Gi complex with the HiBiT system were collected at the Cryo-Electron Microscopy Center of Liangzhu laboratory, Zhejiang University (Hangzhou, China). Imaging was performed using a Titan Krios transmission electron microscope (Thermo Fisher Scientific) operated at 300 kV, equipped with a Gatan K2 Summit direct electron detector.

Automated data collection was carried out using SerialEM software. A total of 6,008 movies were recorded in counting mode with a defocus range of -0.5 to -2.5 μm. Each movie was acquired with a total exposure time of 8 seconds, fractionated into 40 frames at a dose rate of about 8.0 e/Å2/s with a defocus ranging from -0.5 to -2.5 μm using the SerialEM software^38^. The total exposure time was 8 s and 40 frames were recorded per micrograph.

### Image processing and map construction

Motion correction and dose-weighting of the cryo-EM image stacks were performed using MotionCor2.1^39^. The contrast transfer function (CTF) parameters were estimated using Gctf^40^. Particle selection, 2D classification, and 3D classification were carried out using RELION-3.0-beta2^41^.

For BX471-CCR1-Gi complex with LgBiT-HiBiT system, 3,991,512 particles yielded by automated particle picking were subjected to 2D classification, and two rounds of 3D classification using CCL15-CCR1-Gi complex low-pass filtered map as an initial reference model, resulting in a well-defined subset with 482,845 particles^29^. The selected subsets were subsequently subjected to 3D classification with a mask on the receptor or receptor-Gi complex, respectively. High quality particles were selected form the intersection of the best class form these two 3D classifications, producing 205,378 particles. The selected particles were subsequently subjected to 3D refinement, CTF refinement, Bayesian polishing, generating a map with an indicated global resolution of 2.6 Å at a Fourier shell correlation of 0.143. Local resolution was determined using the Bsoft package with half maps as input maps^42^.

For the BX471-bound inactive CCR1, a total of 2,007,636 particles were auto-picked using Laplacian-of-Gaussian in RELION. The particles were then imported into CryoSPARC for iterative rounds of 2D classification and ab-initio reconstruction to generate the initial reference maps, followed by five rounds of heterogeneous refinement in CryoSPARC. The good particles were selected and imported back to RELION for two rounds of 3D classification with soft mask on the complex and mask on the receptor in sequence. A well-defined class of particles was subjected to 3D refinement, CTF refinement and Bayesian polishing. The polished particles were imported into CryoSPARC for further local 3D classification. The high-quality particles for the receptor were processed with local refinement, generating the globally refined map with an indicated resolution of 2.96 Å. The final map was used for subsequent model building and analysis.

For the BX471-bound active CCR1, a total of 2,725,479 particles were auto-picked using Laplacian-of-Gaussian in RELION. These particles were transferred to CryoSPARC for multiple rounds of 2D classification and ab-initio reconstruction to produce the initial reference maps, followed by three rounds of heterogeneous refinement in CryoSPARC. The good particles were selected and imported back to RELION for two rounds of 3D classification with soft mask on the complex and mask on the receptor in sequence. A well-defined class of particles was subjected to 3D refinement, CTF refinement and Bayesian polishing. The polished particles were returned into CryoSPARC for further local 3D classification. The high-quality particles for the receptor were processed with local refinement, generating the globally refined map with an indicated resolution of 2.85 Å. This final map was employed for subsequent model construction and analysis.

### Model building and refinement

The initial CCR1, Gi and scFv16 complex was generated from the CCL15-CCR1-Gi complex (PDB ID: 7VL9)^29^. Then modes were docked into the cryo-EM density map using chimera. After the initial docked models were refined using Rosetta, the models were subjected to iterative rounds of manual adjustment and auto refinement in Coot and Phenix, respectively. The final refinement scores were validated by the module “comprehensive validation (cryo-EM)” in Phenix. Structure figures were prepared by PyMOL, Chimera and ChiemraX.

### Molecular dynamics simulations

The simulation system was derived from the BX471-CCR1-Gi complex. Prior to initiating the simulations, Gβ and Gγ proteins were excluded. The complex was incorporated into a 115×115 Å POPC lipid bilayer using the packmol-memgen software, and was surrounded by a 12 Å aqueous layer^43^. The ionic strength was maintained at 0.15 mol/L NaCl, with additional counterions to balance the system. The FF19SB, Lipid21, and GAFF2 force fields were employed for amino acids, lipids, and ligands, respectively^44–46^. Each system underwent minimization and a heating-equilibration process following established protocols^47,48^. Three independent 500 ns production runs were performed using pmemd.cuda in Amber22 under the NVT ensemble at 300 K and 1 atm^49^. Long-range electrostatic interactions were calculated using the Particle Mesh Ewald method, while a 10 Å cutoff was applied for short-range electrostatic and van der Waals interactions. The SHAKE algorithm was utilized to constrain hydrogen-containing bonds, allowing a timestep of 2 fs. CPPTRAJ was employed to compute RMSD, RMSF, and distances^50^.

### G-protein signaling assay (NanoBiT)

The NanoBiT-G protein dissociation assay was performed as previously described^30^. The NanoBiT-G protein recruitment assay was used to monitor real-time interactions between G protein-coupled receptors (GPCRs) and G proteins. In this system, a small fragment of the NanoLuc Binary Technology (NanoBiT) luciferase (SmBiT) was fused to the C-terminus of the GPCRs, while the large fragment (LgBiT) was inserted into the α-helical domain of the Gα subunit using 15-amino acid flexible linkers. HEK 293T cells (ATCC) were co-transfected with four plasmids using jetPRIME® transfection reagent (Polyplus) according to the manufacturer’s instructions. The plasmids used for transfection were: pcDNA3.1-LgBiT-Gαi (encoding the LgBiT-tagged Gαi subunit), pcDNA3.1-Gβ1 (encoding the Gβ1 subunit), pcDNA3.1-Gγ2 (encoding the Gγ2 subunit), and pcDNA3.1-GPCR-SmBiT (encoding the GPCR fused to SmBiT). The cells were incubated at 37°C with 5% CO2 for 24 hours post-transfection to allow for protein expression. After transfection, the cells were seeded into a 384-well white, opaque plate (Corning) pre-treated with a cell adhesion reagent (Applygen) at a density of 10,000 cells per well. The cells were allowed to adhere for 12 hours at 37°C with 5% CO2. Prior to the assay, the cells were washed three times with Dulbecco’s phosphate-buffered saline (D-PBS) containing 0.5 mM EDTA to remove any residual medium. The cells were then incubated with 25 μL of 5 μM coelenterazine h (Yeasen), a substrate for NanoBiT luciferase, prepared in assay buffer (Hanks’ Balanced Salt Solution (HBSS) with 0.01% (w/v) bovine serum albumin (BSA) and 5 mM HEPES, pH 7.5). Following a 2-hour incubation at room temperature, baseline luminescence was measured using a TECAN multimode microplate reader with an integration time of 500 ns per well. Ligands were prepared in assay buffer at various concentrations and added to the wells in a volume of 5 μL. Immediately after ligand addition, luminescence was recorded to capture the reconstitution of NanoBiT luciferase, reflecting the GPCR-G protein interaction. The raw luminescence data were analyzed using GraphPad Prism 9 software. The signals were normalized against the luminescence values from wells treated with blank ligand (assay buffer) to assess the ligand-induced recruitment of G proteins. Dose-response curves were generated by plotting the normalized luminescence against the logarithm of ligand concentration, and the half-maximal effective concentration (EC50) values were calculated using non-linear regression analysis.

The Ligands used are shown below. The purification method of CCL15 is as described above. Angiotensin II (HY-13948) for AT1R and AT2R, Serotonin(HY-B1473) for 5HT1eR, NPY protein(HY-P71063) for NPY2R,CX3CL1 protein(HY-P7180) for CX3CR1 are from MedChemexpress. CCL17(Cat. No.:C599) for CCR4, IL8(Cat. No.:C035) for CXCR1 and CXCR2, CCL2(Cat. No.:CM78) for CCR2, CCL3(Cat. No.:C061) for CCR5 are from Novoprotein. FMLP(T7091) for FPR1 and FPR2 is from Topscience. CP-55940(CAS No.:83002-04-4) for CB1 is from Sigma.

### Endocytosis measured by flow cytometry

Human monocytic THP1 cells (ATCC-TIB-202) were cultured in RPMI 1640 medium (Hyclone) supplemented with 10% fetal bovine serum (FBS, Invitrogen) and antibiotics (Sangon Biotech). When THP1 cells reached over 80% confluency in a 10 cm dish, they were harvested and plated onto a clear 96-well plate (Corning) at a density of 1 × 10^5 cells per well. The cells were treated with 0.1 μM BX471 (MedChemExpress, Cat. No. HY-12080) for 1 hour at 37°C. Then, the cells were incubated with various concentrations of CCL15 truncation variants (e.g., CCL15(26-92), CCL15(29-92), and CCL15(30-92)) for 2 hours at 37°C to induce receptor endocytosis. Following ligand treatment, the cells were stained with APC-conjugated anti-human CCR1 antibody (Biolegend, Cat #362907) at 4°C to label cell surface receptors. The cells were then counterstained with 4’,6-diamidino-2-phenylindole (DAPI, Sigma-Aldrich) to exclude dead cells during analysis. The samples were analyzed using a CytoFLEX flow cytometer (Beckman Coulter) with the following settings: APC fluorescence was excited at 633 nm and detected using a 660/20 nm bandpass filter, while DAPI fluorescence was excited at 405 nm and detected using a 450/45 nm bandpass filter. Data were acquired and analyzed using CytExpert software (Beckman Coulter). The level of receptor endocytosis was quantified by calculating the normalized endocytosis signal using the following equation: Normalized endocytosis signal = (Median APC fluorescence intensity of cells treated with CCL15 truncation variant) / (Median APC fluorescence intensity of cells treated with the lowest ligand concentration).

#### Acknowledgements

We would like to acknowledge the substantial contribution of Prof. Huahao Shen, who conceived and designed this study. Prof. Huahao Shen passed away in April 2024. He is remembered not only for his professional competence and scientific achievements but also as a man of generous spirit, warm-heartedness, and sense of humor. He sought to build bridges of collaboration, not only within China but also globally with colleagues who shared his passion for improving the care of patients with asthma. His guidance, inspiration, and unwavering support were instrumental in bringing this work to fruition. We are deeply saddened by his loss and dedicate this work to his memory.

This work was supported by grants from the National Natural Science Foundation of China (8230026 to Z.S., 82270023 and U22A20265 to W.L., 32430051, 92353303 and 32141004 to Y.Z., 82225001 and 81920108001 to S.Y.), the Ministry of Science and Technology (2019YFA0508800 to Y.Z), the “Pioneer” and “Leading Goose” R&D Program of Zhejiang (2024C03147 to Y.Z.), the Fundamental Research Funds for the Central Universities (226-2022-00205 to Y.Z.), and National Key Research and Development Program of China (2021YFA1102001 to S.Y.).

## Author contributions

W.L., Y.Z., S.Y. and H.E.X. conceived, designed and supervised the overall project. Z.S. and B.Y. purified the BX471-CCR1-G_i_ complex and prepared the final samples for cryo-EM studies. Z.S. and R.J. purified the inactive BX471-CCR1 complex and prepared the final samples for cryo-EM studies. D.S., C.M. and Q.Y. performed data collection and electron microscopy data processing. Q.S. performed the model building of the BX471-CCR1-G_i_ complex. Y.D. performed the model building of the BX471-CCR1 complex. Z.S., R.J. and B.Y. performed the functional assays with the assistance of C.Z. X.H. performed the molecular dynamics simulation studies. Z.S. and Q.S. wrote the manuscript with guidance of Y.Z., H.E.X., S.Y. and W.L.

## Data availability statement

Cryo-EM maps of BX471-CCR1-Gi and BX471-CCR1 complexes have been deposited in the Electron Microscopy Data Bank under accession codes EMD-XXXXX and EMD-XXXXX, respectively. The atomic coordinates of BX471-CCR1-Gi and BX471-CCR1 complexes have been deposited in the Protein Data Bank under accession codes XXXX and XXXX, respectively. All other data are available upon request to the corresponding authors.

---

**Extended Data Fig. 1|.**
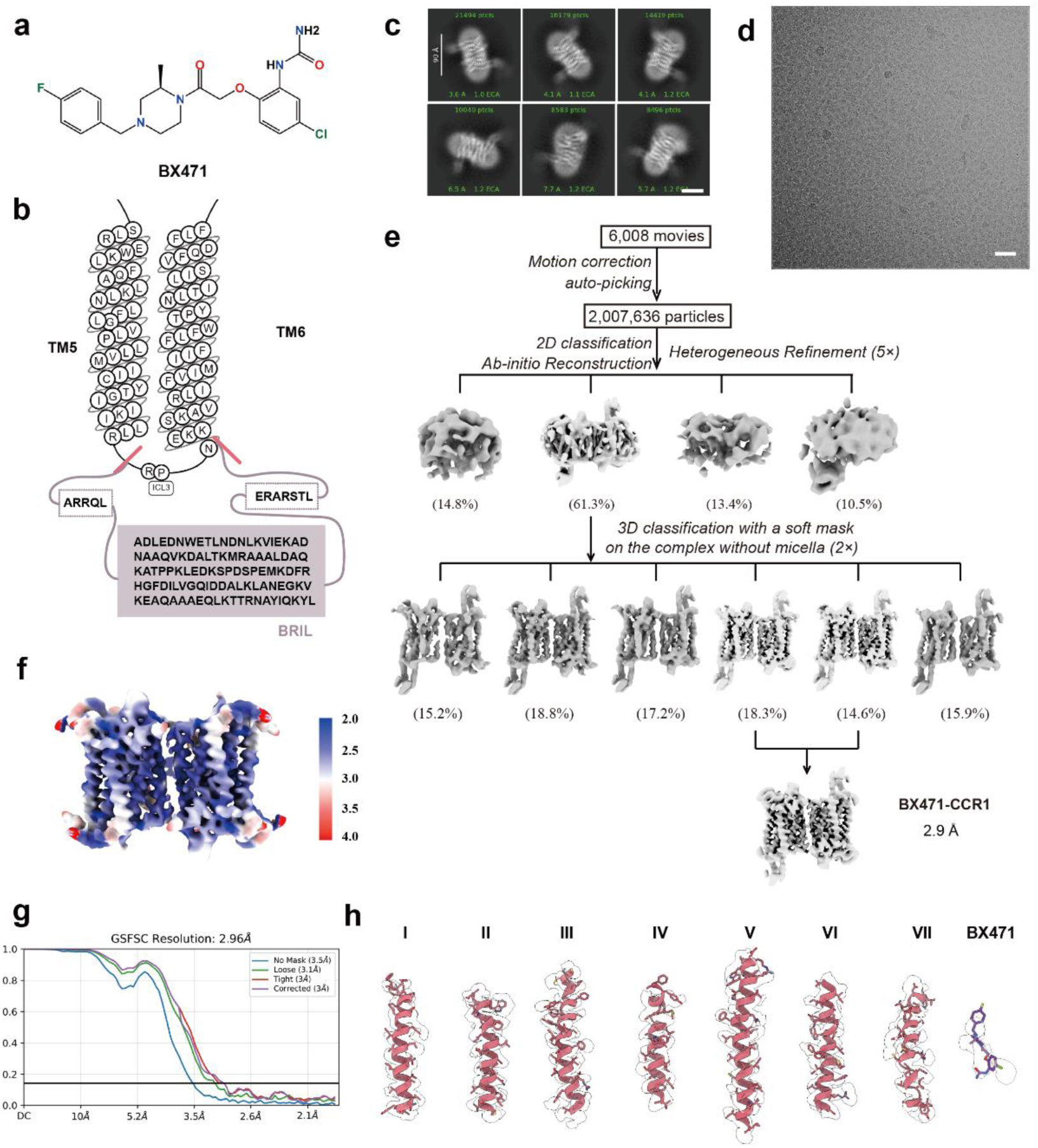
Cryo-EM analysis of the BX471-CCR1 complex. **a**, Chemical structure of BX471. **b,** Detailed illustration of the construct designed for the inactive BX471-CCR1 complex, featuring a BRIL fusion protein to stabilize the receptor. **c,** Representative cryo-EM micrograph of the BX471-CCR1 complex. Scale bar, 50 nm. **d,** Representative 2D class averages showing distinct secondary structure features of the BX471-CCR1 complex from different angles. Scale bar, 5 nm. **e,** Flowchart of cryo-EM data processing and 3D reconstruction for the BX471-CCR1 complex. **f,** Local resolution distribution of the final cryo-EM map of the BX471-CCR1 complex, calculated using the Bsoft package. **g,** Gold-standard FSC curves for the final 3D reconstruction of the BX471-CCR1 complex. **h,** Cryo-EM density map (mesh) and the atomic model (cartoon) of the BX471-CCR1 complex, showing the quality of the map and the fit of the model for all transmembrane helices (TM1-7) and the bound BX471 ligand (purple).

**Extended Data Fig. 2|.**
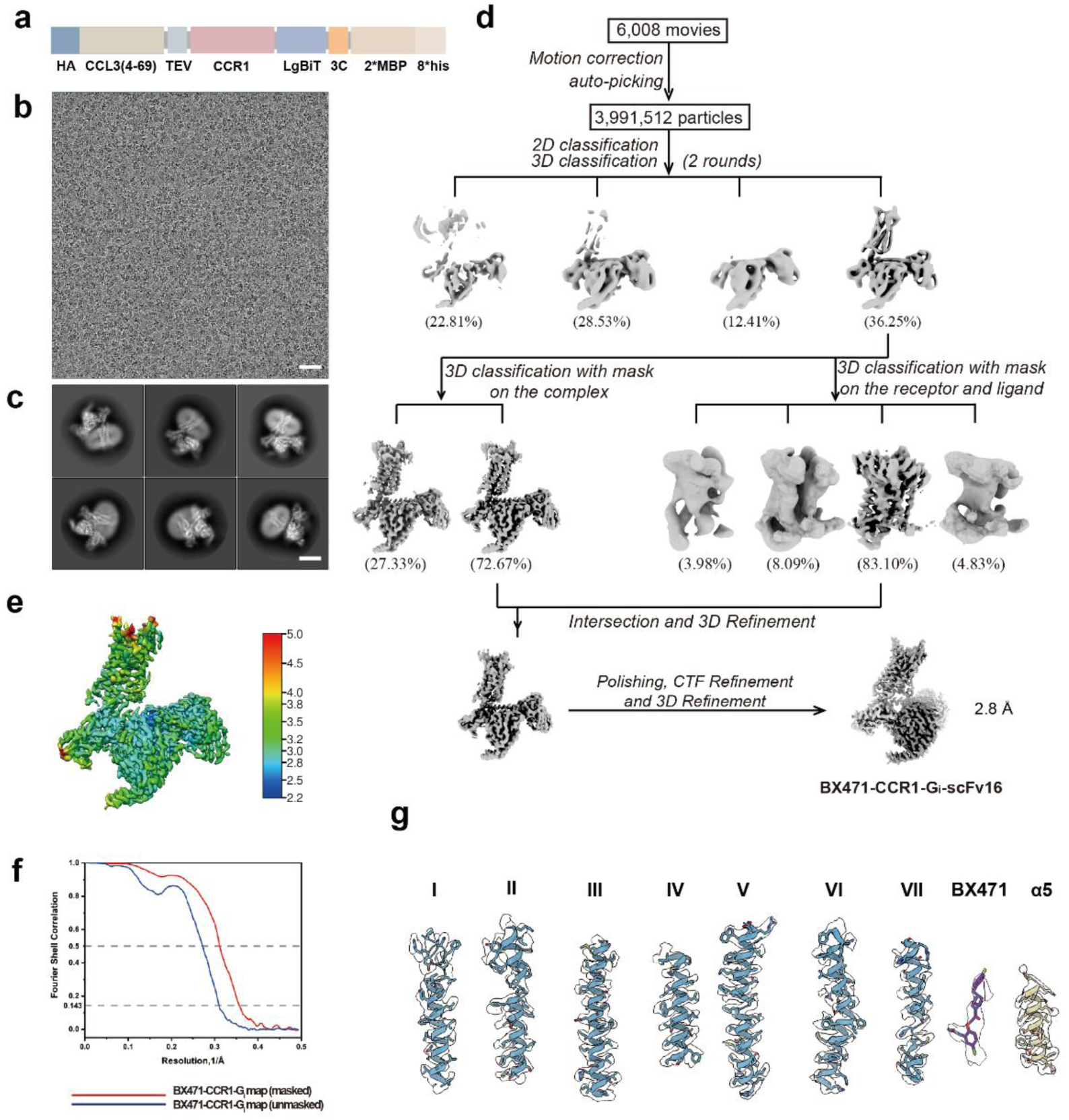
Cryo-EM analysis of the BX471-CCR1-G_i_ complex. **a**, Schematic diagrams of human CCR1-related constructs used in this study for the structural determination of the BX471-CCR1-Gi complex. The constructs include wild-type CCR1 with a C-terminal SmBiT tag, and the engineered Gαi1 with an inserted LgBiT tag and an N-terminal 8×His tag for purification. **b,** Representative cryo-EM micrograph of the BX471-CCR1-Gi complex. Scale bar, 30 nm. **c,** Representative 2D class averages showing distinct secondary structure features of the BX471-CCR1-Gi complex from different angles. Scale bar, 5 nm. **d,** Flowchart of cryo-EM data processing and 3D reconstruction for the BX471-CCR1-Gi complex. **e,** Local resolution distribution of the final cryo-EM map of the BX471-CCR1-Gi complex, calculated using the Bsoft package. **f,** Gold-standard FSC curves for the final 3D reconstruction of the BX471-CCR1-Gi complex. **g,** Cryo-EM density map (mesh) and the atomic model (cartoon) of the BX471-CCR1-Gi complex, showing the quality of the map and the fit of the model for all transmembrane helices (TM1-7), the bound BX471 ligand (purple), and the α5-helix of the Gαi subunit (yellow).

**Extended Data Fig. 3|.**
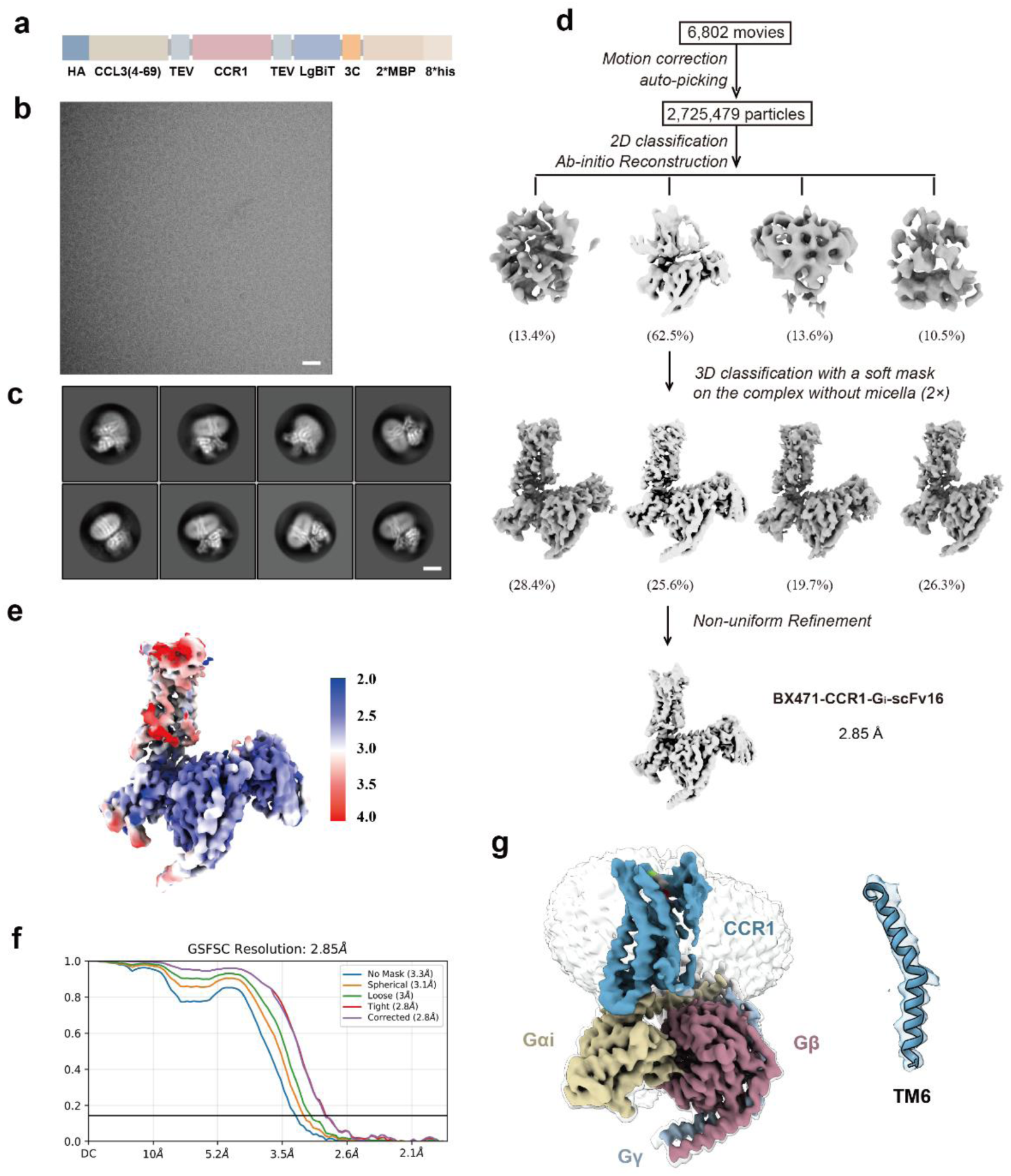
Cryo-EM analysis of the BX471-CCR1-G_i_ complex in the absence of LgBiT-HiBiT system. **a,** Schematic diagrams of human CCR1-related constructs used in this study for the structural determination of the BX471-CCR1-Gi complex without the LgBiT-HiBiT system. The constructs include CCR1 with a C-terminal 3C protease cleavage site followed by the LgBiT tag and a TEV protease cleavage site, and the engineered Gαi1 with an N-terminal 8×His tag for purification. **b,** Representative cryo-EM micrograph of the BX471-CCR1-Gi complex in the absence of the LgBiT-HiBiT system. Scale bar, 30 nm. **c,** Representative 2D class averages showing distinct secondary structure features of the BX471-CCR1-Gi complex from different angles. Scale bar, 5 nm. **d,** Flowchart of cryo-EM data processing and 3D reconstruction for the BX471-CCR1-Gi complex without the LgBiT-HiBiT system. **e,** Local resolution distribution of the final cryo-EM map of the BX471-CCR1-Gi complex, calculated using the Bsoft package. **f,** Gold-standard FSC curves for the final 3D reconstruction of the BX471-CCR1-Gi complex. **g,** (Left) Cryo-EM density map of the BX471-CCR1-Gi complex in the absence of the LgBiT-HiBiT system, with CCR1 in the SO state (dodger blue), Gαi1 (yellow), Gβ (rosy brown), and Gγ (light blue). (Right) A fit of the TM6 model from the BX471-CCR1-Gi complex with the LgBiT-HiBiT system (cartoon) to the electron density map of the complex without the LgBiT-HiBiT system (mesh), demonstrating the similarity of the TM6 conformation in both structures.

**Extended Data Fig. 4 |.**
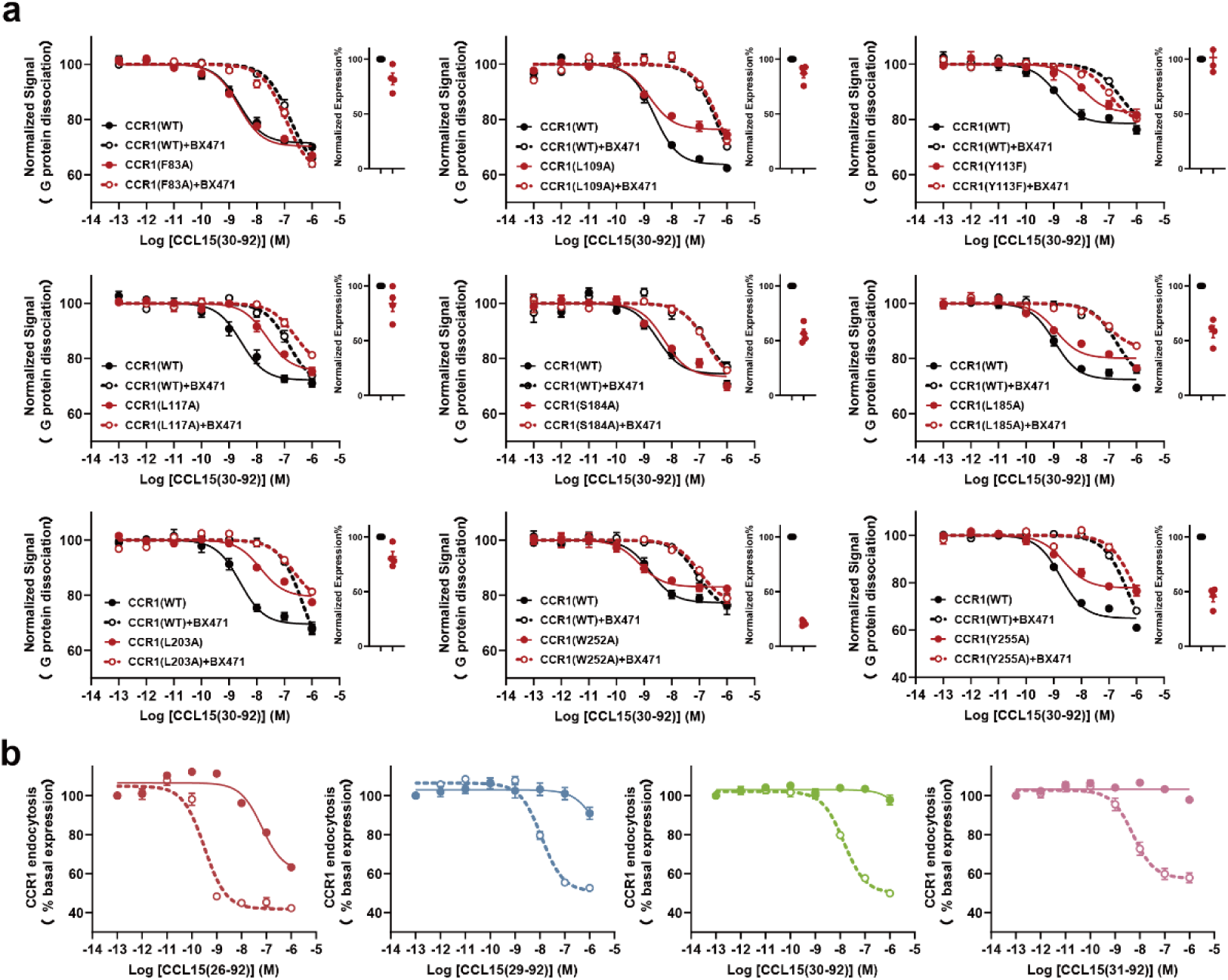
The influence of CCR1 mutations on the inhibitory effect of BX471 on CCL15(30-92)-induced G protein activation and the effect of BX471 on CCL15-induced CCR1 internalization. **a**, Dose-response curves for CCL15(30-92)-induced Gi signaling on wild-type CCR1 and various CCR1 mutants (F83A, L109A, Y113F, Y117A, S184A, L185A, L203A, W252A, and Y255A) in the presence of BX471, measured by the NanoBiT assay. **b**, Dose-response curves for the effect of various CCL15 N-terminal truncation variants, including CCL15(26-92), CCL15(29-92), CCL15(30-92), and CCL15(31-92), on the cell surface expression of CCR1 in the presence of BX471 (1 μM), measured by flow cytometry in THP-1 cells. Data points and error bars represent the mean and SEM, respectively, from *N* = 4 (a) or 5-6 (b) independent experiments performed with single replicates.

**Extended Data Fig. 5|.**
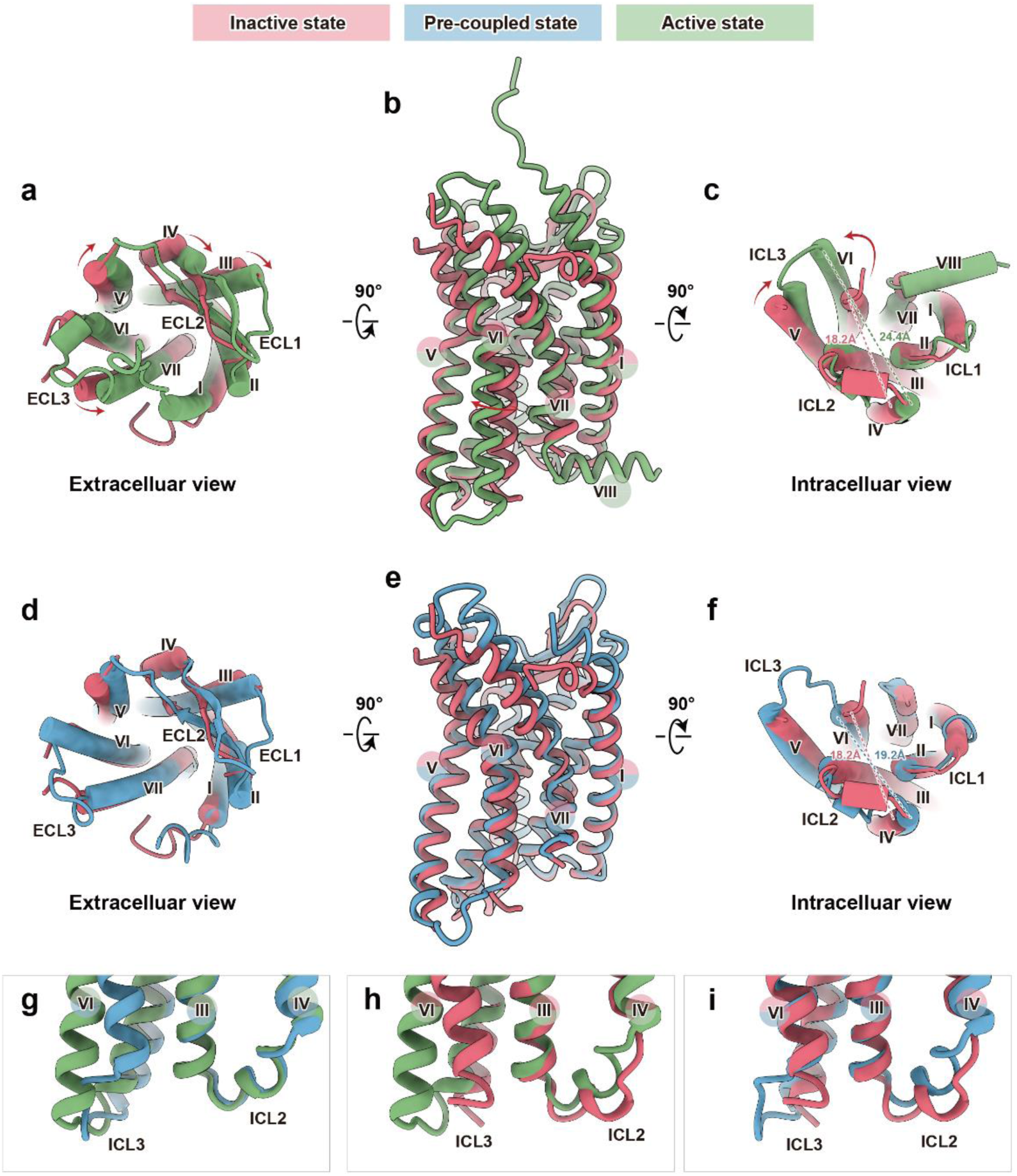
Structural comparison of CCR1 in different states. **a-c**, Superimposed structures of active CCR1 (green, PDB ID: 7VL9) and inactive CCR1 (rose red). **d-f,** Superimposed structures of pre-coupled state CCR1 (dodger blue) and inactive CCR1 (rose red). Extracellular (**a, d**), side (**b, e**), and intracellular (**c, f**) views of the overall structures are presented. **g-i,** Detailed views of the TM3-TM4 region, ICL2, TM6, and ICL3 in pre-coupled (dodger blue) and active (green) CCR1 structures (**g**), inactive (rose red) and active (green) CCR1 structures (**h**), and inactive (rose red) and pre-coupled (blue) CCR1 structures (**i**).

**Extended Data Fig. 6|.**
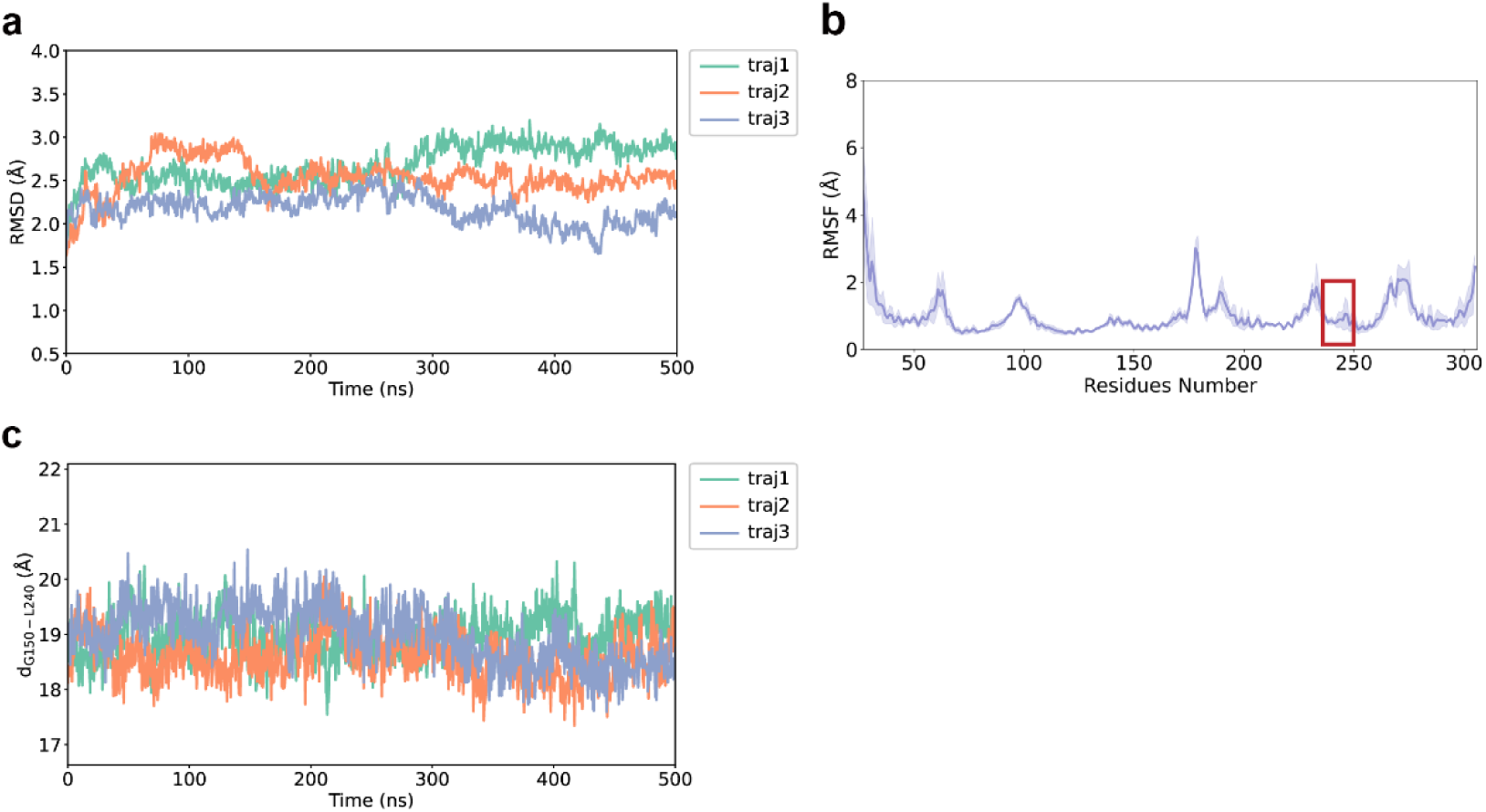
Molecular dynamics simulations of CCR1 in the pre-coupled state. **a**, The root mean standard deviation (RMSD) of the CCR1 structure was stable across all 500 ns production runs, with an average value of 2.46 ± 0.29 Å. **b,** The root mean standard fluctuation (RMSF) data indicate that the most flexible regions are the N-terminus and ECL2 (residues Y170^4.62^-L192^5.31^). The intracellular portion of TM6 (residues K236^6.32^ to L250^6.46^) is highlighted by the red rectangle, with an average RMSF of 1.69 ± 0.32 Å. **c,** The distance between G150^4.42^ and L240^6.46^ across three trajectories.

**Extended Data Fig. 7|.**
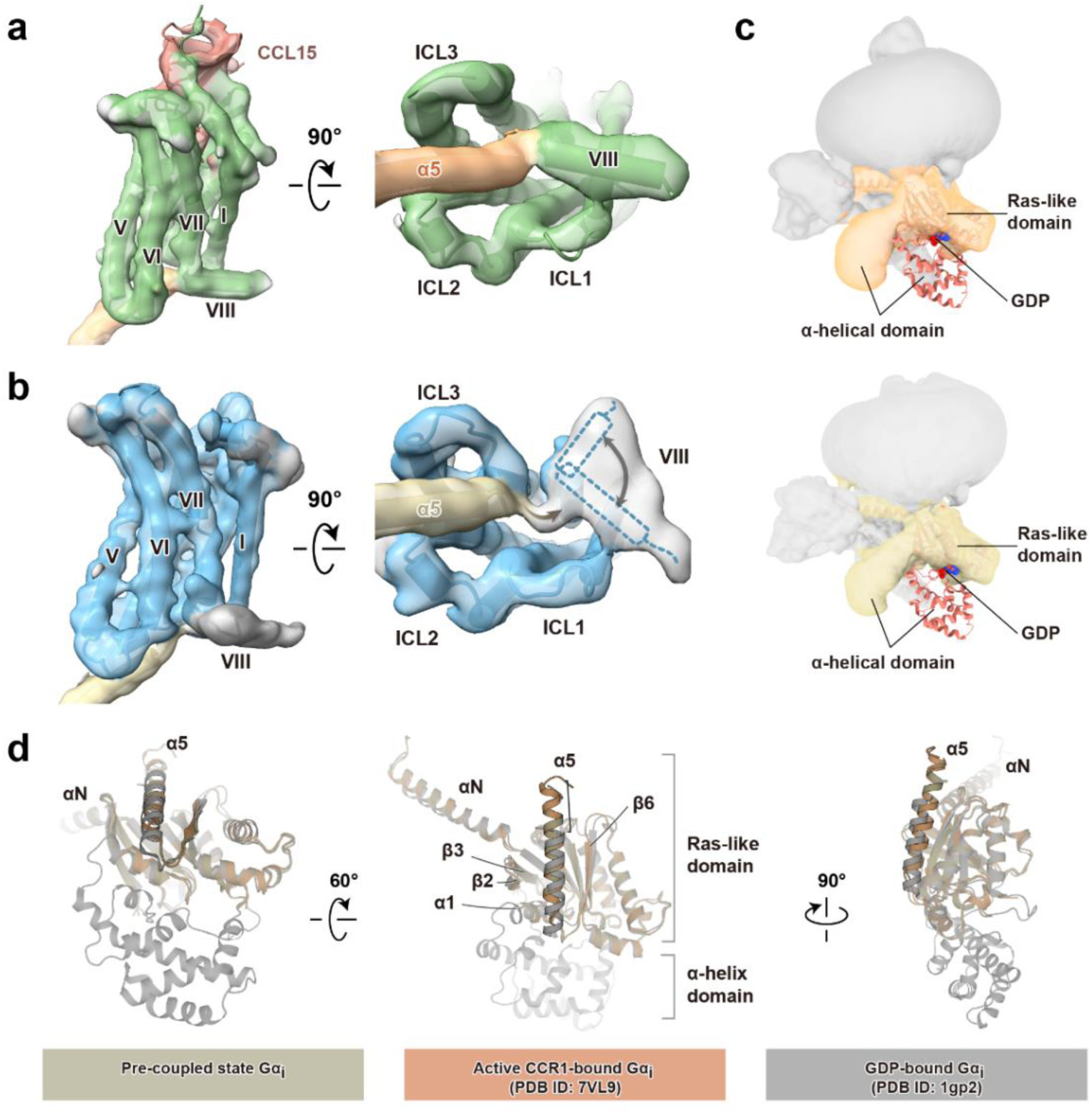
Density maps and models of CCR1-Gi in the active and pre-coupled states. **a-b**, Side (left) and intracellular (right) views of cryo-EM density maps of the CCL15-CCR1-Gi1 (**a,** PDB ID: 7VL9) and BX471-CCR1-Gi1 (**b**) complexes. CCR1 is shown in green (active state) and dodger blue (pre-coupled state), while Gαi1 is shown in gold (active state) and yellow (pre-coupled state). CCL15 is depicted in brown. In **b,** the supposed location of helix 8 (H8) is indicated with a dashed line, based on the observed density. The brown arrow highlights the flexibility of the C-terminus of Gαi1 in the pre-coupled state. **c,** Overlay of the crystal structure of GDP-bound Gαi (orange, PDB ID: 1gp2) with the cryo-EM density maps of Gαi from the CCL15-CCR1-Gi1 complex (gold, upper panel) and the BX471-CCR1-Gi1 complex (yellow, lower panel). GDP is shown as spheres. **d,** Structural alignment of GDP-bound Gαi (grey, PDB ID: 1gp2) with nucleotide-free Gαi1 from the CCL15-CCR1-Gi1 (tan) and BX471-CCR1-Gi1 (dark khaki) complexes.

**Extended Data Fig. 8|.**
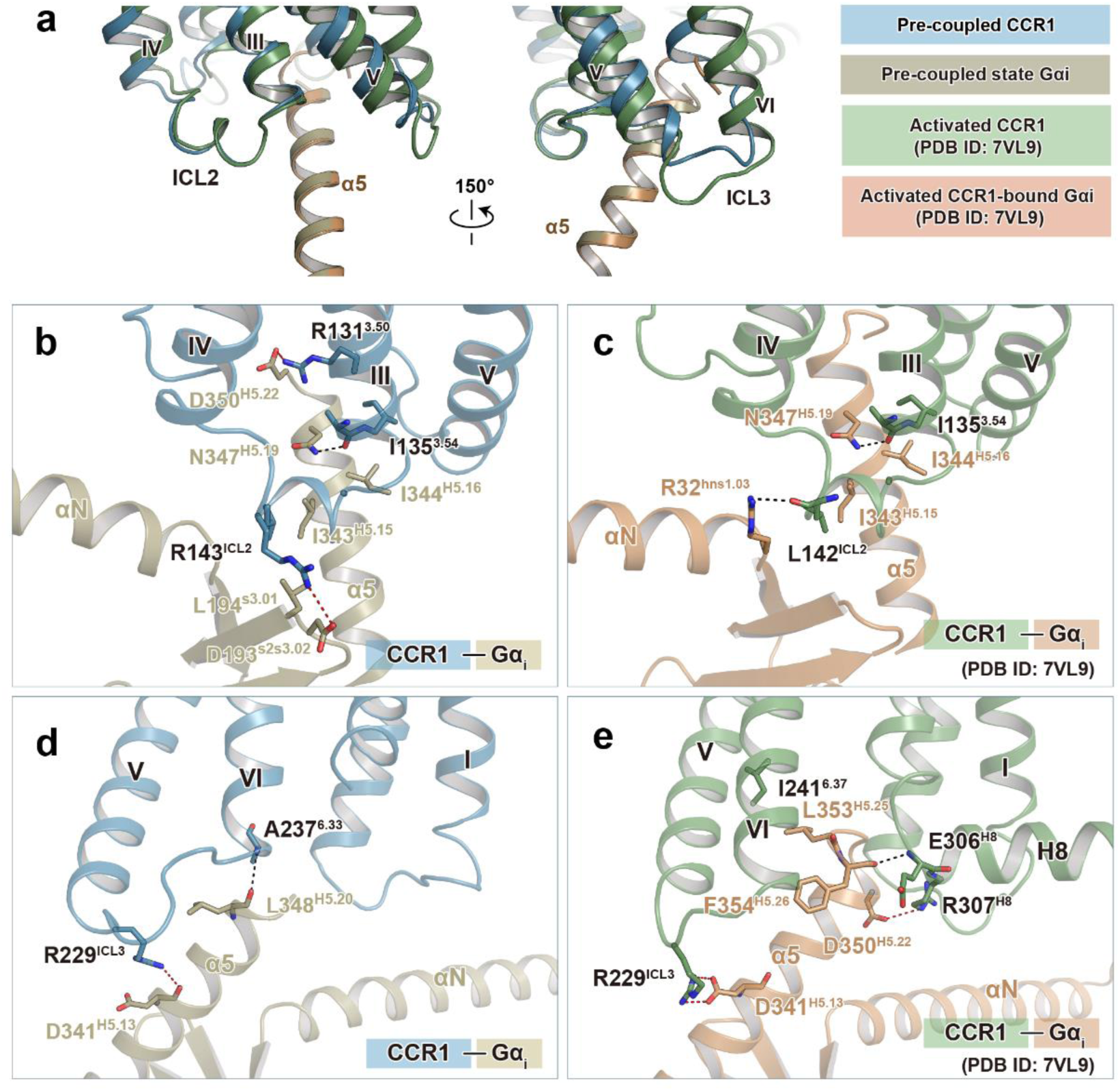
Interactions between CCR1 and Gαi in the active and pre-coupled states. **a**, Comparison of the interface between the α5 helix of Gαi and the cytoplasmic region of CCR1 in the active (green) and pre-coupled (dodger blue) states. The alignment is performed using the Gα protein as the reference. **b-c,** Detailed interactions around ICL2 region between CCR1 (dodger blue) and the Gαi1 protein (yellow) in the pre-coupled (**b**) and active (**c**, PDB ID: 7VL9) states. **d-e,** Detailed interactions around ICL3 region between CCR1 (dodger blue) and the Gαi1 protein (yellow) in the pre-coupled (**d**) and active (**e**, PDB ID: 7VL9) states. Hydrogen bonds are depicted as black dashed lines, while salt bridges are represented by red dashed lines.

**Extended Data Fig. 9|.**
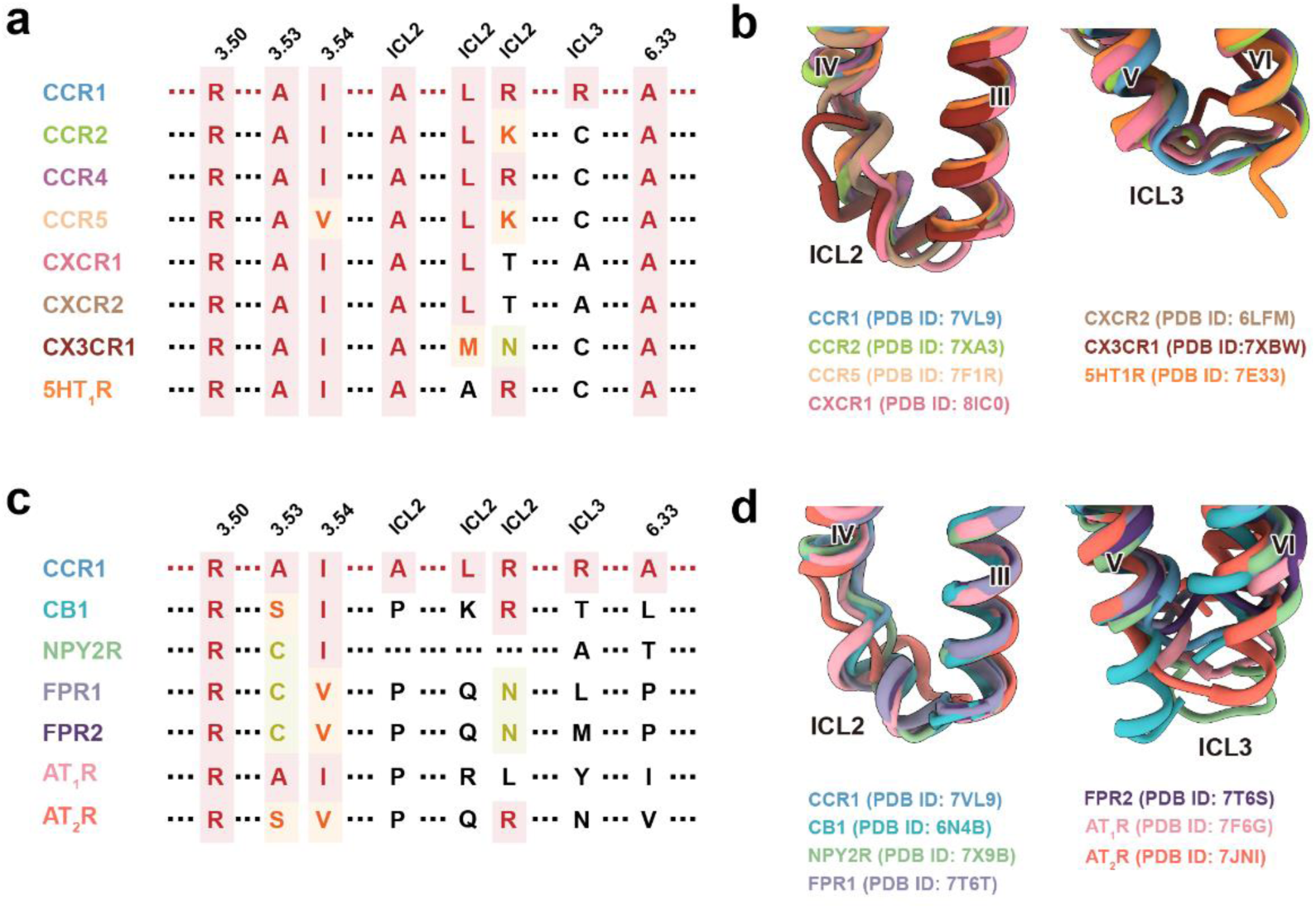
Structural comparison of CCR1 with other GPCRs and a proposed model of G protein coupling. **a-b**, Structure-based sequence alignment (**a**) and comparison of the ICL2 (left panel) and ICL3 (right panel) regions (**b**) among CCR1 and seven other GPCRs with high sequence similarity. **c-d**, Structure-based sequence alignment (**c**) and comparison of the ICL2 (upper panel) and ICL3 (lower panel) regions (**d**) among CCR1 and six other GPCRs with low sequence similarity, with the same color coding as in panel a. In **a** and **c**, Residues with fully similar properties are marked in red, residues with strongly similar properties (scoring > 0.5 in the Gonnet PAM 250 matrix) are marked in orange, and residues with weakly similar properties (scoring < 0.5 in the Gonnet PAM 250 matrix) are marked in yellow.

**Supplementary Table 1.** | Cryo-EM data collection, model refinement and validation statistics.

|  | BX471-CCR1-Gi<br>(with LgBiT-HiBiT) | BX471-CCR1-Gi | BX471-CCR1 |
| --- | --- | --- | --- |
| **Data collection and processing** |  |  |  |
| Magnification | 81,000 | 150,540 | 150,540 |
| Voltage (kV) | 300 | 300 | 300 |
| Electron exposure (e<sup>-</sup>/Å<sup>2</sup>) | 70 | 51 | 51 |
| Defocus range (μm) | -0.5 ~ -3.0 | -1.0 ~ -2.0 | -1.0 ~ -2.0 |
| Pixel size (Å) | 1.071 | 0.93 | 0.93 |
| Symmetry imposed | C1 | C1 | C1 |
| Initial particle projections (no.) | 5,812,667 | 2,007,636 | 2,007,636 |
| Final particle projections (no.) | 1,090,180 | 305,378 | 196,117 |
| Map resolution (Å) | 2.6 | 2.9 | 3.0 |
| FSC threshold | 0.143 | 0.143 | 0.143 |
| Map resolution range (Å) | 2.5-5.0 | 2.0-4.0 | 2.0-4.0 |
| **Refinement** |  |  |  |
| Initial model used | 6DO1 | 6DO1 | AlphaFold2 |
| Model resolution (Å) | 2.7 |  | 3.0 |
| FSC threshold | 0.5 |  | 0.5 |
| Map sharpening *B* factor (Å<sup>2</sup>) | -89.73 |  | -121.20 |
| **Model composition** |  |  |  |
| Non-hydrogen atoms | 9605 |  | 2068 |
| Protein residues | 1214 |  | 250 |
| Lipid | 0 |  | 1 |
| Water | 1 |  | 0 |
| ***B* factors (Å<sup>2</sup>)** |  |  |  |
| Protein | 46.54 |  | 30.37 |
| Lipids | 39.97 |  | 75.28 |
|  |  |  | \ |
| **R.m.s. deviations** |  |  |  |
| Bond lengths (Å) | 0.007 |  | 0.004 |
| Bond angles (°) | 0.937 |  | 0.811 |
| **Validation** |  |  |  |
| MolProbity score | 1.60 |  | 2.82 |
| Clashscore | 8.10 |  | 8.73 |
| Rotamer outliers (%) | 0.00 |  |  |
| **Ramachandran plot** |  |  |  |
| Favored (%) | 97.07 |  | 93.70 |
| Allowed (%) | 2.93 |  | 6.30 |
| Disallowed (%) | 0 |  | 0 |

